# Cyclic loading regime considered beneficial does not protect injured and interleukin-1-inflamed cartilage from post-traumatic osteoarthritis

**DOI:** 10.1101/2021.08.23.457262

**Authors:** ASA Eskelinen, C Florea, P Tanska, HK Hung, EH Frank, S Mikkonen, P Nieminen, P Julkunen, AJ Grodzinsky, RK Korhonen

## Abstract

Post-traumatic osteoarthritis is a degenerative musculoskeletal condition where homeostasis of articular cartilage is perturbated by lesions and inflammation, leading to abnormal tissue-level loading. These mechanisms have rarely been included simultaneously in *in vitro* osteoarthritis models. We modeled the early disease progression in bovine cartilage regulated by the coaction of **(1)** mechanical injury, **(2)** pro-inflammatory interleukin-1α challenge, and **(3)** cyclic loading mimicking walking and considered beneficial (15% strain, 1 Hz). Surprisingly, cyclic loading did not protect cartilage from accelerated glycosaminoglycan loss over 12 days of interleukin-1-culture despite promoting aggrecan biosynthesis. Our time-dependent data suggest that this loading regime could be beneficial in the first days following injury but later turn detrimental in interleukin-1-inflamed cartilage. Consequently, early anti-catabolic drug intervention may inhibit, whereas cyclic loading during chronic inflammation may promote osteoarthritis progression. Our data on the early stages of post-traumatic osteoarthritis could be utilized in the development of countermeasures for disease progression.

## 1 Introduction

Osteoarthritis is the most prevalent musculoskeletal disorder after low back pain and fractures with hundreds of millions of patients suffering from pain and disability^1^. One of the hallmarks of this disease is the degeneration of articular cartilage. The injury-related disease phenotype — post-traumatic osteoarthritis (PTOA) — manifests itself in the knee joint as the presence of chondral lesions^2^, altered mechanical loading environment^3^, and elevated whole-joint inflammatory response^4^. Despite the vast body of literature, the mechanobiological response of cartilage to simultaneous mechanical damage, pro-inflammatory cytokine infiltration, specifically interleukin-1 (IL-1), and physiological cyclic loading remain to be elucidated. Here, we utilized an experimental platform^5^ to study these mechanisms concomitantly for the first time with IL-1 in order to gain crucial insights into the complex and multifaceted progression of early-stage PTOA. The obtained time-dependent and cartilage depth-dependent data can be useful for the development of methods and treatments tackling against disease progression.

The response of cartilage to injurious loading, inflammation, and cyclic loading have been separately studied in the past few decades. First, injurious loading has been shown to lead to accelerated loss of sulfated glycosaminoglycans (GAG; found for example attached to aggrecan, the major proteoglycan in cartilage) in the extracellular matrix (ECM)^6–10^, localized especially near lesions^11^. Biosynthesis of new aggrecan with native GAG chains is not significantly altered in the superficial nor in the deeper cartilage^8^ but biosynthesis of aggrecan with abnormal GAG (alterations in the sulfation pattern of new chondroitin sulfate chains) may increase in deeper tissue shortly after injury^12^ or intermittently applied cyclic compression^13^. Injurious loading is also suggested to cause marked chondrocyte (cartilage cell) death^12, 14^ eminently in the vicinity of lesions^11^ and to increase oxidative stress^15^, which in turn might further promote apoptosis^16, 17^. Second, inflammation induced by pro-catabolic cytokines such as IL-1, IL-6, and tumor necrosis factor-α (TNFα) results in an accelerated depletion of GAG from cartilage^5, 18, 19^ in a dose-dependent manner^10, 19^ via increased expression of proteolytic enzymes^20, 21^. Furthermore, inflammatory agents elicit a marked immune response in chondrocytes; the biosynthesis of aggrecan decreases and cell death increases^5, 18^. Third, cyclic loading is suggested to decrease^11, 22^ or increase^23^ the GAG content depending on the aggressiveness of loading. Aggressiveness of cyclic loading is specified by several mechanical variables, such as loading amplitude, frequency, waveform, and duration, which have been given a wide variety of values and configurations in the literature^24^. Generally, moderate cyclic loading (*e.g.*, 10–20% strain amplitude, 1 Hz loading frequency, applied continuously for up to 48h or intermittently in three or four 1h periods per day for up to 14 days) is widely considered beneficial for cartilage health, as demonstrated by the increased biosynthesis of new proteoglycans^5, 24–26^. Moderate cyclic loading may also mitigate chondrocyte apoptosis^27^ compared to more aggressive loading (such as 30% strain amplitude which was considered overloading and resulting in pro-inflammatory cartilage response)^5^. Although physiological cyclic loading could be within healthy and normal limits at the level of the whole joint, the tissue-level loading and cartilage deformation could be abnormal locally and thus lead to localized GAG loss or apoptosis^28^. Additionally, physiological mechanical loading appears to stimulate cells to release IL-4^29^, which has potent anti-inflammatory and anti-catabolic roles in the suppression of the degradative effects of IL-1^30^ and TNFα^31^. Such beneficial cyclic loading (10 and 20% strain amplitudes applied with 0.5 Hz frequency, 1h on/5h off per day) has been shown to effectively mitigate GAG loss after 4 and 8 days even when antagonized by exogenous IL-6 and TNFα^5^. Similar protective effect of cyclic loading has been observed also with the presence of exogenous IL-1, but this has been studied only over 1–3 days without initial controlled injurious loading of cartilage^32–34^. Without cyclic loading, an injured tissue environment predisposes cartilage to faster inflammation-affected degeneration compared to mechanically intact environment^5, 35^.

The purpose of this *in vitro* study with bovine cartilage was to determine the effects of simultaneous injurious loading, IL-1-culture and moderate cyclic loading on time-dependent bulk and localized GAG loss, aggrecan biosynthesis, and cell viability beyond the early response up to 12 days (Fig. 1). We aimed to fill a void in previous literature regarding the behavior of IL-1 in an injured and cyclically loaded environment; the same has been previously studied with cytokines IL-6 and TNFα^5^. In the light of the previous studies^5, 11, 12, 18, 32–35^, we hypothesized that a combination of injurious loading and IL-1 challenge would be more degrading than either of these conditions alone, but our chosen cyclic loading regime (15% strain amplitude, 1 Hz, haversine waveform, 1h on/5h off per day and including an overnight rest; see Supplementary Material S1, Section S1, Fig. S1), hypothesized to be beneficial based on earlier literature^5, 24–26^, would ameliorate the degradation. Contrary to our hypothesis, we observed that moderate cyclic loading did not inhibit or inhibited only mildly the pro-catabolic response of cartilage to relatively high concentration (1 ng/ml) of IL-1 over 12 days. Even though cyclic loading increased the biosynthesis of aggrecan and occasionally led to locally high GAG levels away from lesions especially in the deep tissue, the data suggest that cyclic loading applied with our protocol cannot rescue the bulk cartilage from degradation during IL-1-mediated inflammation. Interestingly, we also noted an increased depletion of GAG soon after injurious loading and introduction of IL-1, when cyclic loading was not present, but this early degradation was suppressed when cyclic loading was involved. Overall, our data suggest that a cyclic loading regime previously thought to be advantageous may be beneficial only initially after joint trauma but may become an adverse factor over prolonged time periods in case the inflammatory response is not removed.

**Fig. 1.**
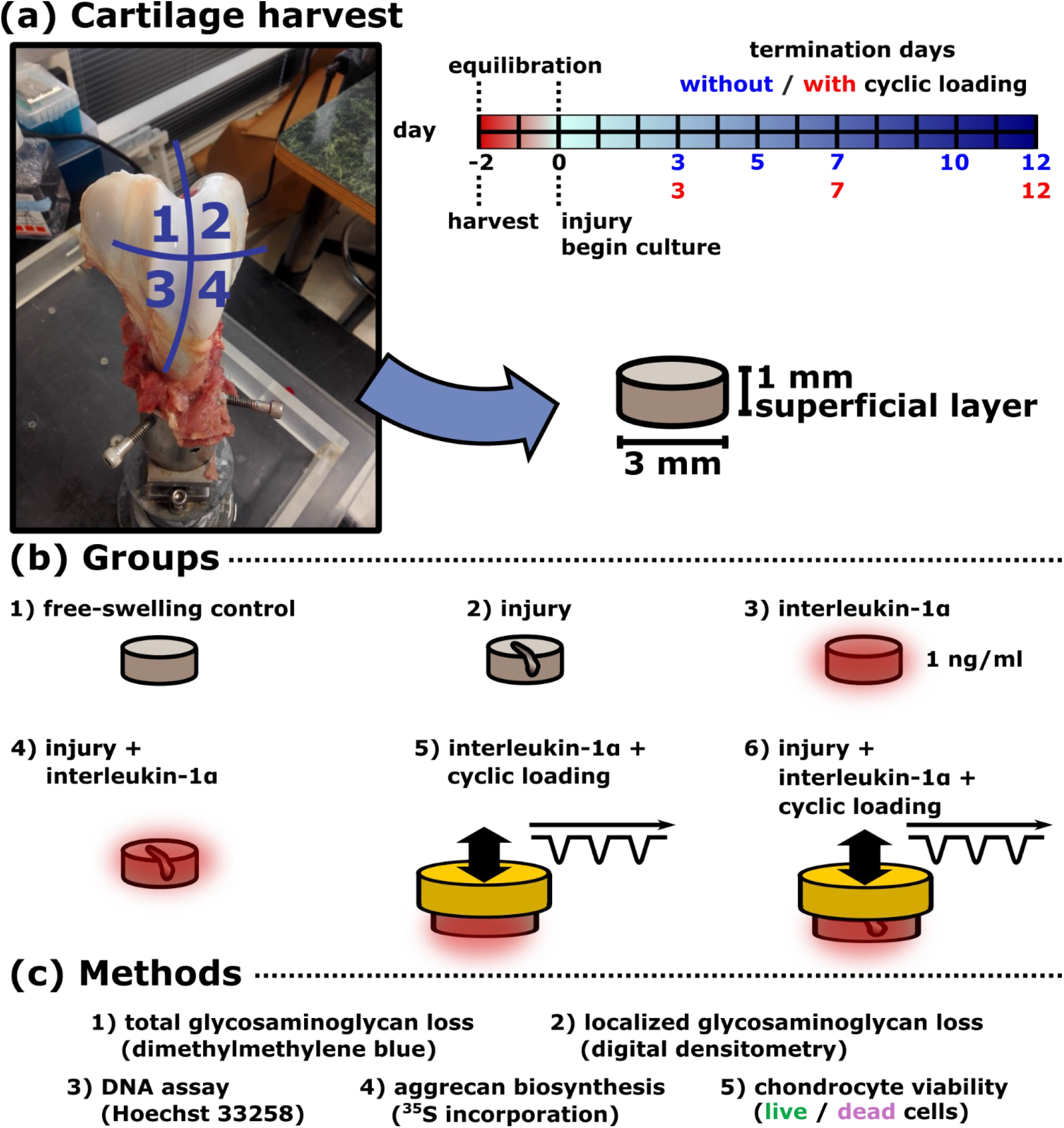
Cartilage tissue harvest, treatment groups and used methods. **(a)** Cartilage plugs were collected from the patellofemoral grooves of 1-2-week-old calves (*N* = 6 animals). The surface of each groove was divided into four areas, all of which provided samples to **1)** treatment groups without cyclic loading and **2)** cyclically loaded groups as well as to animal and location-matched free-swelling control groups. After a 2-day equilibrium period, the samples were subjected to **(b)** six different culture conditions for up to 12 days: **1)** free-swelling serving as control (CTRL); groups without cyclic loading were **2)** injury-only (INJ), **3)** exogenous interleukin-1α-challenge-only (IL), and **4)** injured and IL-1α-challenged (INJ-IL); the cyclically loaded groups were **5)** IL-1α-challenged and cyclically loaded (IL-CL) and **6)** injured, IL-1α-challenged, and cyclically loaded (INJ-IL-CL). **(c)** To determine the cartilage response to the different culture conditions, we used glycosaminoglycan assay of the whole plugs (in total *n* = 6 location-matched plugs from two different animals per treatment and analysis time point), optical density measurements indicative of localized glycosaminoglycan (GAG) loss (*n* = 6), and DNA (*n* = 6), aggrecan biosynthesis (*n* = 6) and cell viability (*n* = 2) assays.

## 2 Results

### 2.1 IL-1α increases GAG release in a dose-dependent manner

In order to determine a suitable concentration of IL-1α for the rest of the study, we first conducted 7-day and 12-day cultures of calf cartilage plugs with only the biochemical degradation mechanism present in the form of exogenous cytokine challenge (free-swelling as control (CTRL), 0.1, 0.5, 1, and 10 ng/ml of IL-1α). Accumulated GAG loss (%) — a measure of GAG release over time from a cartilage plug in relation to the total GAG content of that plug (Eq. 1; see Methods Subsection 4.3) — was higher and faster with increasing cytokine concentration. From day 4 onwards, 1 ng/ml of IL-1 caused significantly more accumulated GAG loss compared to CTRL conditions; for example, on day 12, the average accumulated GAG losses in 1 ng/ml and CTRL groups were 21.7% and 15.7%, respectively (*p* = 0.003; linear mixed effects model, detailed statistics are compiled into Supplementary Material S1). The concentration of 1 ng/ml of IL-1α was chosen for the rest of the study since this concentration was able to cause a substantial and fast degradation compared to the CTRL conditions. This concentration was also low enough to not to be overwhelmingly destructive for the ECM (like 10 ng/ml would likely have been, based on the pilot study) and, thus, to not dominate over biomechanical degradative factors (injury, cyclic loading). For more details, see the Supplementary Material S1 (Subsections S2.1-2.2, Figs. S2 and S3, Tables S1-S6).

### 2.2 Cyclic loading decreases GAG loss of IL-1α-affected cartilage initially but its inhibitory effects are hampered after 12 days of loading

We subjected the cartilage plugs to injurious loading, exogenous cytokine challenge (1 ng/ml of IL-1α), and cyclic loading (for the cyclic loading protocol, see Supplementary Material S1, Section S1, Fig. S1). The cartilage plugs were assigned into the following treatment groups: **1)** free-swelling control (referred as CTRL in the text), **2)** injury-only (INJ), **3)** IL-1α-challenge-only (IL), **4)** injured and IL-1α-challenged (INJ-IL), **5)** IL-1α-challenged and cyclically loaded (IL-CL), and **6)** injured, IL-1α-challenged, and cyclically loaded (INJ-IL-CL; Fig. 1b).

The accumulated GAG loss was the highest in the INJ-IL group throughout the experiment (Fig. 2). During the first 2–4 days after the beginning of the treatment, the accumulated GAG loss in the INJ-IL group (for example, day 4: average 18.4%) was significantly higher than in the CTRL (11.8%, *p* < 0.001), INJ (15.7%, *p* = 0.003) and IL groups (15.6%, *p* = 0.002). The early 2–4-day GAG release response was lower in cyclically loaded groups IL-CL (for example, day 4: average 12.3%) and INJ-IL-CL (12.6%) than in the INJ-IL group, although the differences were non-significant (IL-CL *vs.* INJ-IL *p* = 0.247; INJ-IL-CL *vs.* INJ-IL *p* = 0.378). After 8 days, the degradative effects of cytokines became pronounced; the IL group with 25.8% of average accumulated GAG loss showed more GAG depletion than the INJ group (22.6%, *p* = 0.018). On day 12, the INJ-IL group (average 40.0%) exhibited significantly higher accumulated GAG loss than its corresponding cyclically loaded group INJ-IL-CL (29.8%, *p* = 0.037). In uninjured inflamed plugs, cyclic loading decreased accumulated GAG loss, but this difference was non-significant (IL average 33.1% *vs.* IL-CL 25.7%, *p* = 0.877). See more details in the Supplementary Material S1 (Subsection S2.3, Tables S7-S9).

**Fig 2.**
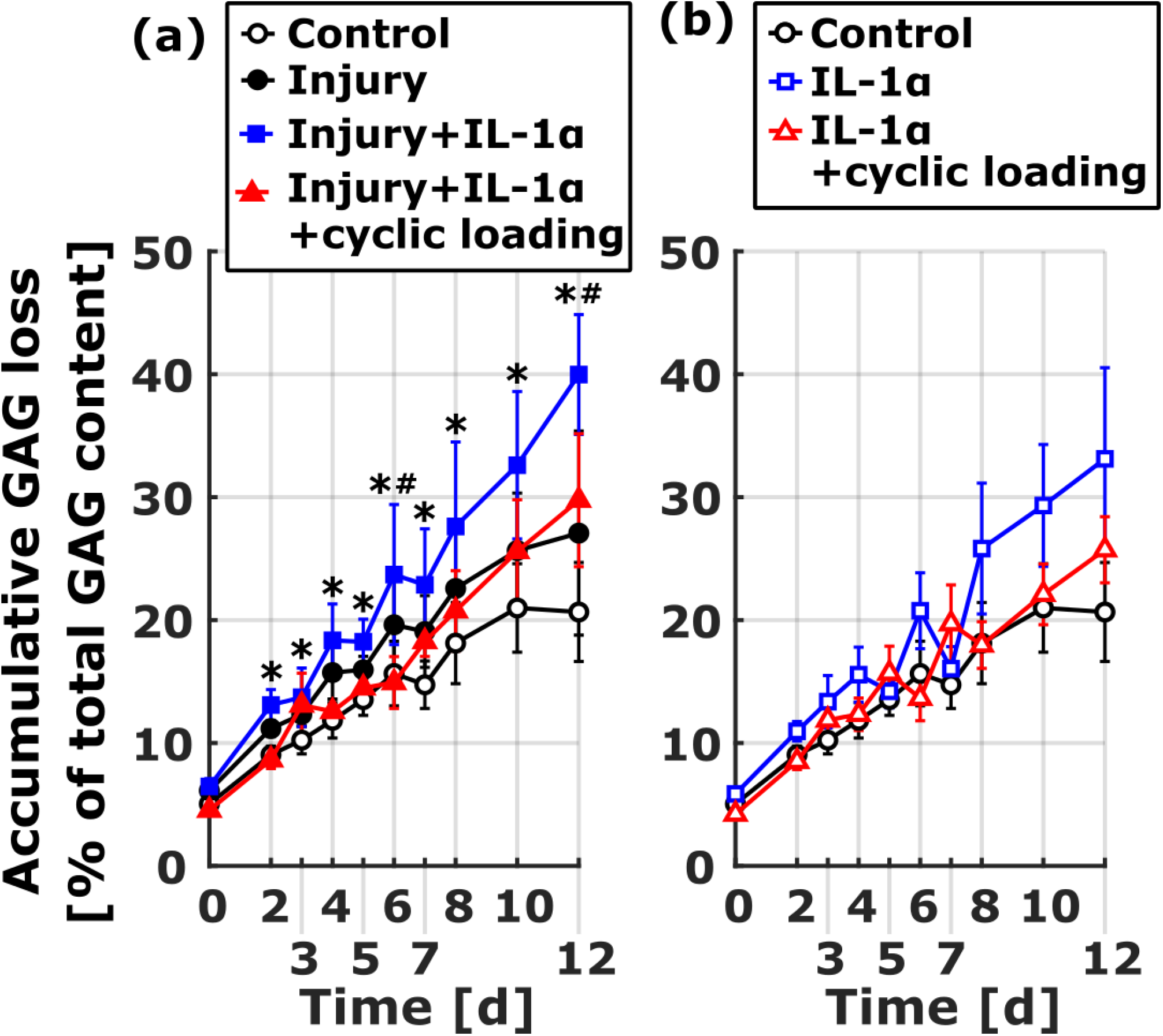
Accumulated glycosaminoglycan (GAG) loss. **(a)** The INJ-IL group exhibited the greatest accumulated GAG loss that was significantly higher compared to the INJ group from day 2 onwards (data shown as mean ± 95% confidence intervals). On day 12, the accumulated GAG loss in the INJ-IL group was also significantly higher than in the INJ-IL-CL group (*p* = 0.037). **(b)** Also, the IL group showed more accumulated GAG loss than the IL-CL group on day 12, but this difference was non-significant (*p* = 0.877). Our accumulated GAG loss data suggest that cyclic loading might decrease GAG loss in injured and IL-1-challenged cartilage, but in general it does not protect or the proteoglycan matrix protection is only mild for IL-1-challenged cartilage. See also Fig. 3 for normalized accumulated GAG loss data suggesting that cyclic loading could even be deleterious for inflamed cartilage. The data was collected at different time points from different animals, leading to apparent decrease of mean accumulated GAG loss between time points 3–4, 5–6, and 7–8 days; however, individual plugs exhibited strictly increasing accumulated GAG loss over time. Linear mixed effects model, *p* < 0.05: * = INJ-IL *vs.* INJ, # = INJ-IL *vs.* INJ-IL-CL. Abbreviations: INJ = injury-only, IL = IL-1α-challenge-only, INJ-IL = injured and IL-1α-challenged, IL-CL = IL-1α-challenged and cyclically loaded, INJ-IL-CL = injured, IL-1α-challenged, and cyclically loaded.

Next, we present the data as normalized accumulated GAG loss. The normalization was calculated as a fold change between accumulated GAG loss (% of total) of a treated plug compared to that of an animal- and location-matched CTRL plug (Eq. 2; see Methods Subsection 4.3). This allows comparison of proteoglycan matrix damage in response to different treatments without a bias from animal- or cartilage location-related differences in GAG depletion. The value of 1 means that the treated plugs accumulated their GAG loss (% of total) in the same extent as the control plugs.

The INJ and IL groups showed a significant early (days 2–4 from the beginning of treatment) increase in the normalized accumulated GAG loss (Fig. 3ab). For example, between days 0 and 3, the normalized accumulated GAG loss increased from a median fold change of 1.02 to 1.25 in the INJ group (*p* < 0.001) and from 1.01 to 1.42 in the IL group (*p* < 0.001). By day 12, the GAG loss fold changes decreased to a median of 1.08 in the INJ group (day 12 *vs.* day 3, *p* = 0.111) and to 1.33 in the IL group (*p* = 0.737). The INJ-IL group exhibited a steadily increasing normalized accumulated GAG loss, reaching ultimately a median of 1.66 on day 12 (significantly higher than on day 0: 1.17, *p* < 0.001; Fig. 3c). In turn, cyclically loaded samples (Fig. 3de) did not show significant early increase in normalized accumulated GAG loss in contrast to samples without cyclic loading (Fig. 3a-c). For example, between days 0 and 3, the normalized accumulated GAG loss increased from a median of 1.08 to 1.15 in the IL-CL group (*p* = 0.197) and decreased from 1.21 to 1.16 in the INJ-IL-CL group (*p* = 0.091). However, the cyclically loaded groups exhibited significant increase in normalized accumulated GAG loss until day 12 in both IL-CL (median 1.47 on day 12; day 12 *vs*. day 3 *p* = 0.001; Fig. 3d), and INJ-IL-CL groups (1.69 on day 12; day 12 *vs*. day 3 *p* < 0.001; Fig. 3e). See more details in the Supplementary Material S1 (Subsection S2.3, Fig. S4, Tables S10-S12).

**Fig. 3.**
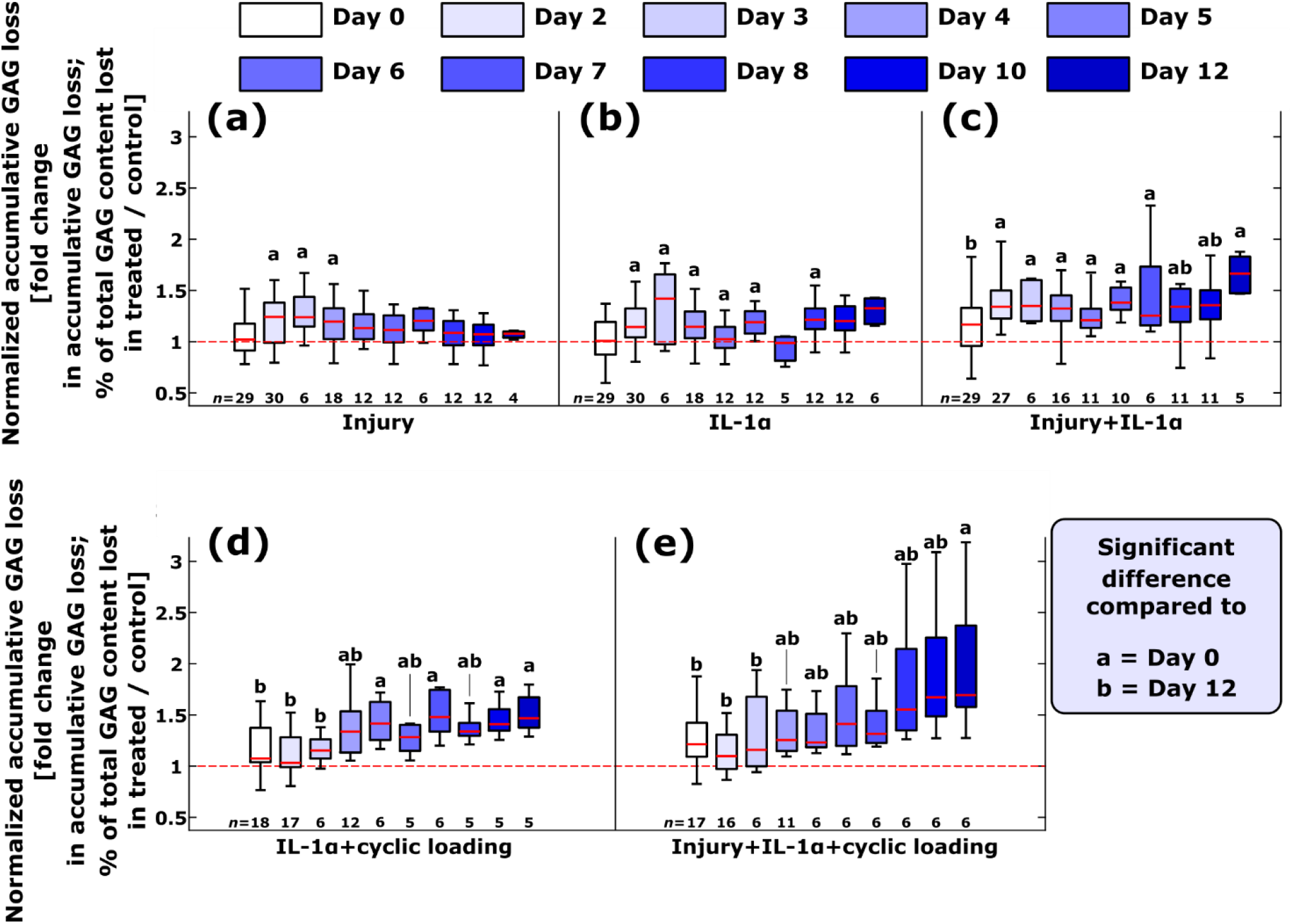
Normalized accumulated glycosaminoglycan (GAG) loss in different treatment groups at different time points. The accumulated GAG loss in **(a)** injury-only (INJ), **(b)** interleukin-1α-challenge-only (IL), **(c)** injured and IL-1α-challenged (INJ-IL), **(d)** IL-1α-challenged and cyclically loaded (IL-CL), and **(e)** injured, IL-1α-challenged, and cyclically loaded (INJ-IL-CL) plugs was normalized with that of animal and location-matched free-swelling controls. Groups without cyclic loading showed early (2–4 days after beginning of the treatment) increase in normalized accumulated GAG loss while cyclically loaded groups exhibited mitigation of early degenerative response. Normalized accumulated GAG loss increased significantly over the 12-day time period in all except the INJ and IL groups. On day 12, the INJ-IL-CL group exhibited significantly higher normalized accumulated GAG loss compared to the INJ-IL group (*p* = 0.007), hinting that cyclic loading does not protect injured and IL-1-inflamed cartilage from GAG loss. The amount of plugs *n* designated for each time point is shown on the bottom of each subpanel. In boxplots, the bars indicate medians, the boxes interquartile ranges (*i.e.*, from 25% to 75% percentiles), and the whiskers minimum and maximum values. The letters above the boxplots indicate statistically significant differences in normalized accumulated GAG loss between time points of 0, 2, 8, and 12 days after initiation of the treatment (*p* < 0.05; linear mixed effects model).

Starting from day 2 after the initiation of the treatment, the INJ-IL group exhibited significantly higher normalized accumulated GAG loss compared to the INJ and IL groups (S1 Supplementary Material, Subsection S2.3, Fig. S4). On day 2, the normalized accumulated GAG loss of the INJ-IL group (median 1.34) was higher than in the INJ (median 1.23, *p* = 0.003), IL (1.14, *p* = 0.001), and also in the cyclically loaded IL-CL (1.03, *p* = 0.019) and INJ-IL-CL groups (1.10, *p* = 0.044). On day 12, the INJ-IL-CL group (median 1.69) exhibited the highest levels of normalized accumulated GAG loss and was significantly higher than in the INJ-IL (median 1.66, *p* = 0.007) and IL-CL groups (1.47, *p* = 0.031; Fig. S4j). Moreover, the IL-CL group had higher normalized accumulated GAG loss (median 1.47) than the IL group on day 12 (1.32, *p* = 0.005).

### 2.3 Cyclic loading protects the deep regions of inflamed cartilage from GAG loss following three days of treatment

In general, qualitative optical density maps (low optical density is indicative of GAG loss) of Safranin-O-stained cartilage sections showed extensive GAG loss close to the chondral lesions (Fig. 4). Plugs treated with IL-1 showed gradual decrease of localized GAG concentration also in intact regions away from lesions, as well as by the lateral edges.

**Fig. 4.**
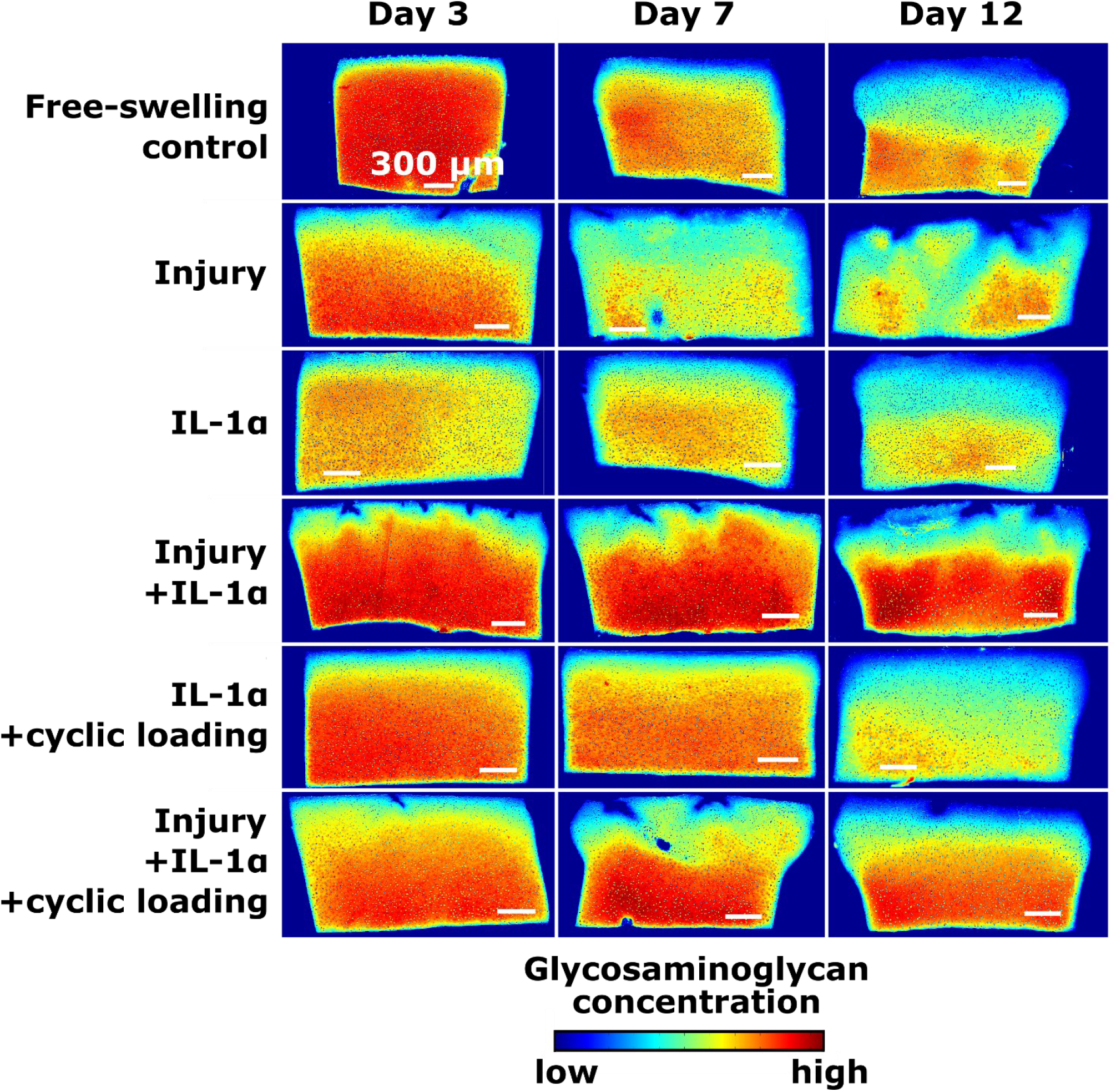
Localized glycosaminoglycan (GAG) content in qualitative optical density maps of Safranin-O-stained sections. Red and blue areas of high and low optical density are indicative of high and low localized GAG content, respectively. In general, GAG content decreased substantially locally near lesions whereas intact regions did not exhibit as intensive localized GAG loss. Interleukin (IL)-1α-cultured plugs showed gradual decrease in localized GAG content starting from the surface and lateral edges of the plug, leaving the central and deep tissue regions relatively rich in GAG concentration. Each treatment and time point combination shown here had six plugs prepared for digital densitometry analysis (three slices per plug from the middle of the plug).

Next, optical density profiles were taken from the optical density maps. Based on depth-wise optical density profiles taken away from lesions after three days from the beginning of the treatment (Fig. 5a; S1 Supplementary Material, Subsection S2.4, Fig. S5a), the INJ and IL groups experienced significantly more localized GAG loss through the whole plug depth compared to the CTRL group (*p* < 0.05 throughout Section 2.3.). The INJ-IL group retained GAG levels away from lesions close to control levels. In cyclically loaded groups, the depth-dependent GAG content was below that of CTRL group. Specifically, the GAG content in the IL-CL group decreased especially in the superficial and transitional zones, but was retained in the deeper matrix at similar or higher levels as in the INJ and IL groups. Moreover, the optical density in the lower transitional and deep zones (>40% of the normalized depth) of the INJ-IL-CL group was significantly higher than in the INJ and IL groups (Figs. 5a and S5a). On day 7 (Fig. 5b), the INJ and INJ-IL-CL groups depleted their depth-wise GAG away from injuries more than the CTRL group. The depth-wise GAG content was higher in the IL-CL group compared to the CTRL group (Fig. S5b). On day 12 (S1 Supplementary Material, Subsection S2.4, Fig. S6d), none of the treatment group profiles deviated significantly from the CTRL profiles. However, the optical densities of the INJ-IL-CL group were higher than in the INJ-IL group in the deep cartilage regions. See more details in the Supplementary Material S1 (Subsection S2.4, Figs. S5-S7).

**Fig. 5.**
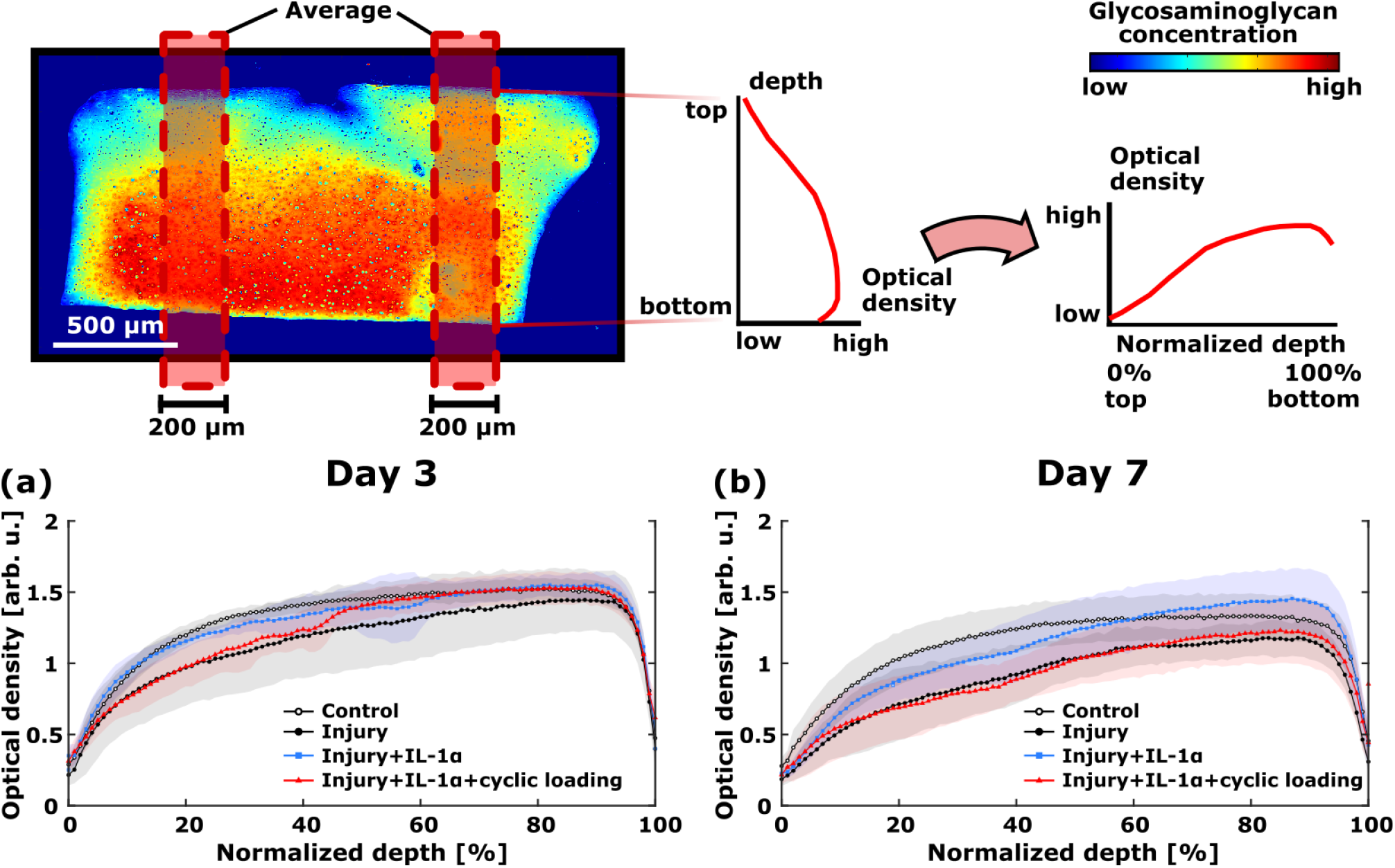
Optical density profiles reveal depth-wise changes in localized glycosaminoglycan (GAG) content in response to different treatments. Two profiles were acquired and averaged from each histological slice (three slices per cartilage plug, six plugs per treatment and time point combination). The profiles were taken from intact areas on both sides of lesions (injured samples) or at the center and middle between center and lateral edge (uninjured samples). **(a)** On day 3, the GAG content decreased significantly (*p* < 0.05; linear mixed effects model; data shown as mean ± standard deviation) in the INJ and IL groups through the whole depth compared to the CTRL group (for the IL and IL-CL group profiles and normalized depths where statistically significant differences were observed, see Supplementary Material, Subsection S2.4, Fig. S5). Cyclically loaded groups depleted GAG especially in the superficial and transitional zones but retained GAG concentration close to or higher than the INJ and IL groups in the deeper tissues. For example, the INJ-IL-CL group had significantly higher optical densities in the lower transitional and deep zones (normalized depths >40%) compared to the INJ group. **(b)** On day 7, the injured groups in general were more deprived of their GAG content compared to uninjured groups. Furthermore, cyclic loading led to more severe GAG loss in the INJ-IL-CL group compared to the INJ-IL group. Abbreviations: CTRL = free-swelling control, INJ = injury-only, IL = IL-1α-challenge-only, INJ-IL = injured and IL-1α-challenged, INJ-IL-CL = injured, IL-1α-challenged, and cyclically loaded.

Next, we assessed localized average optical density in superficial regions near and away from lesions. At all time points, the GAG content diminished significantly in the vicinity of cartilage cracks compared to intact regions away from lesions (*p* < 0.05, Fig. 6a, Supplementary Material S1, Subsection S2.4, Figs. S8ac and S9aceg). On day 3 (Fig. 6b), both INJ-IL and INJ-IL-CL groups exhibited higher regional average GAG concentrations away from lesions compared to intact regions in INJ, IL, and IL-CL groups. On day 7 (Fig. 6c), the average optical density in the intact regions of the IL-CL group was still near the CTRL group levels and higher than the IL group levels. Similarly, on day 12 (Supplementary Material S1, Subsection S2.4, Fig. S9h), the superficial zone showed less localized GAG loss in intact regions of cyclically loaded groups compared to the INJ-IL group.

**Fig. 6.**
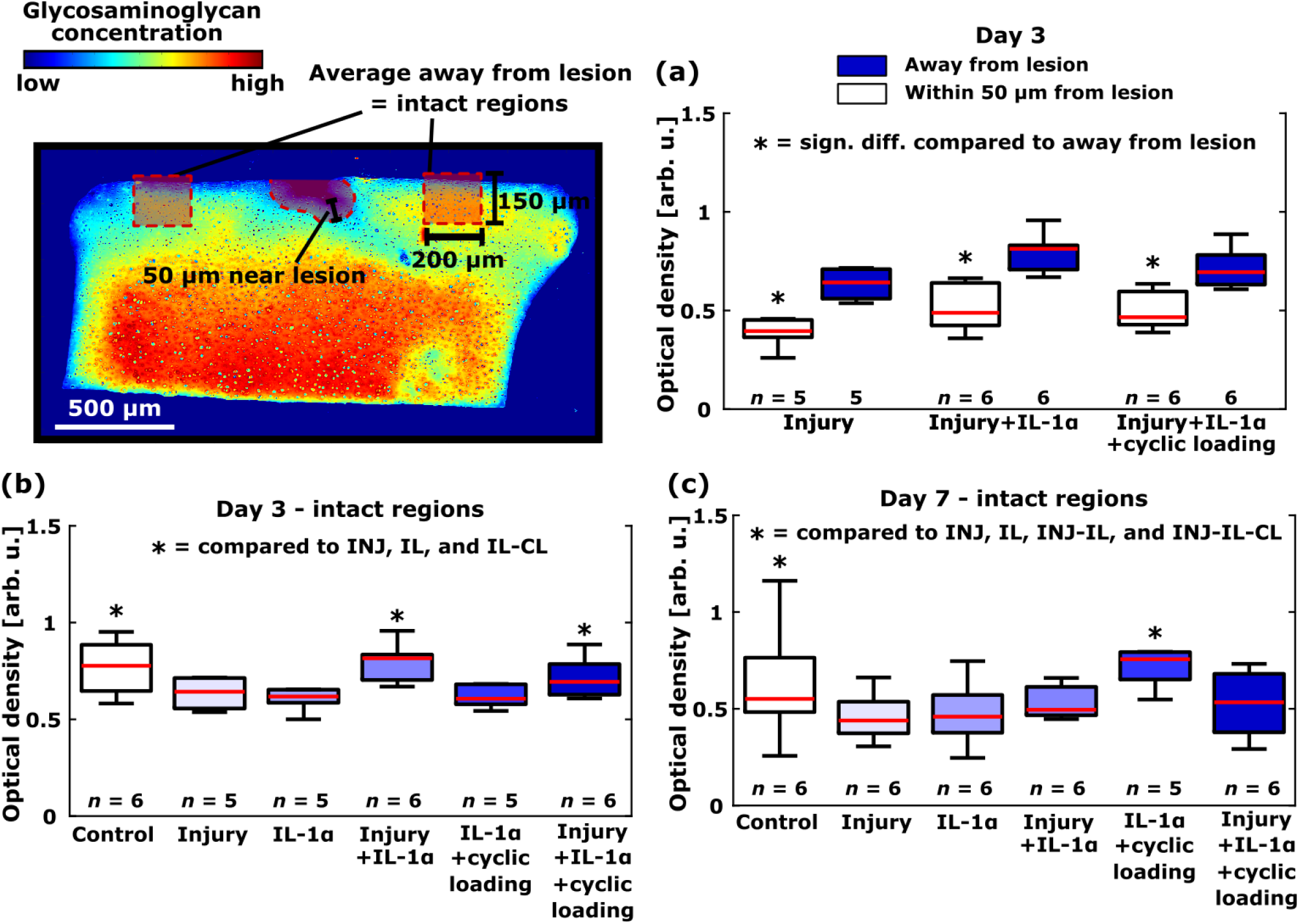
Localized glycosaminoglycan (GAG) content near and away from lesions. One region near and two regions away from lesions were obtained and their average regional optical density was calculated from each histological slice (three slices per cartilage plug, six plugs per treatment and time point combination). The intact areas were taken from both sides of lesions (injured samples) or at the center and middle between center and lateral edge (uninjured samples). On day 3 after beginning of the treatment, **(a)** the localized average GAG content was significantly lower (*p* < 0.05; linear mixed effects model) near the lesions compared to away from lesions in all the injured groups, and **(b)** surprisingly, the INJ-IL and INJ-IL-CL groups retained their GAG content better in regions away from lesions compared to the INJ, IL, and IL-CL groups. On day 7, **(c)** all the other groups except the IL-CL group lost their GAG content away from lesions below that of free-swelling control levels. For more details, see Supplementary Material S1, Subsection S2.4, Fig. S8. The amount of plugs *n* designated for each group is shown on the bottom of subpanels. Abbreviations: INJ = injury-only, IL = IL-1α-challenge-only, INJ-IL = injured and IL-1α-challenged, IL-CL = IL-1α-challenged and cyclically loaded, INJ-IL-CL = injured, IL-1α-challenged, and cyclically loaded.

### 2.4 IL-1α-challenge inhibits aggrecan biosynthesis but moderate cyclic loading can induce aggrecan production even in an injured and inflamed tissue environment

Cytokine challenge as well as CTRL conditions suppressed significantly the aggrecan biosynthesis rate measured with ^35^S incorporation and normalized with the amount of DNA within the cartilage samples between days 3 and 12 (Fig. 7ac). Specifically, in the IL group, the median biosynthesis rate decreased from 68 pmol/h/μg (day 3) to 43 pmol/h/μg (day 12, *p* < 0.001; Fig. 7c). In the INJ group, the aggrecan biosynthesis rate increased on days 7 and 10 (median synthesis rates 96 pmol/h/μg and 128 pmol/h/μg, respectively) compared to day 5 (33 pmol/h/μg; day 7 *vs.* day 5 *p* = 0.028; day 10 *vs.* day 5 *p* = 0.002), but decreased once again close to the day 3–5 levels on day 12 (53 pmol/h/μg; day 10 *vs.* day 12 *p* < 0.001; Fig. 7b). In turn, the INJ-IL group did not show any increase or decrease (day 5 median synthesis rate 30 pmol/h/μg, day 10 rate 40 pmol/h/μg, day 12 rate 44 pmol/h/μg; day 5 *vs.* day 10 *p* = 0.208, day 10 *vs.* day 12 *p* = 0.860). With cyclically loaded groups, there was an increase in aggrecan biosynthesis in both the IL-CL (day 7 median synthesis rate 49 pmol/h/μg, day 12 rate 92 pmol/h/μg, *p* = 0.008; Fig. 7e) and INJ-IL-CL groups (day 7 rate 59 pmol/h/μg, day 12 rate 118 pmol/h/μg, *p* < 0.001).

**Fig. 7.**
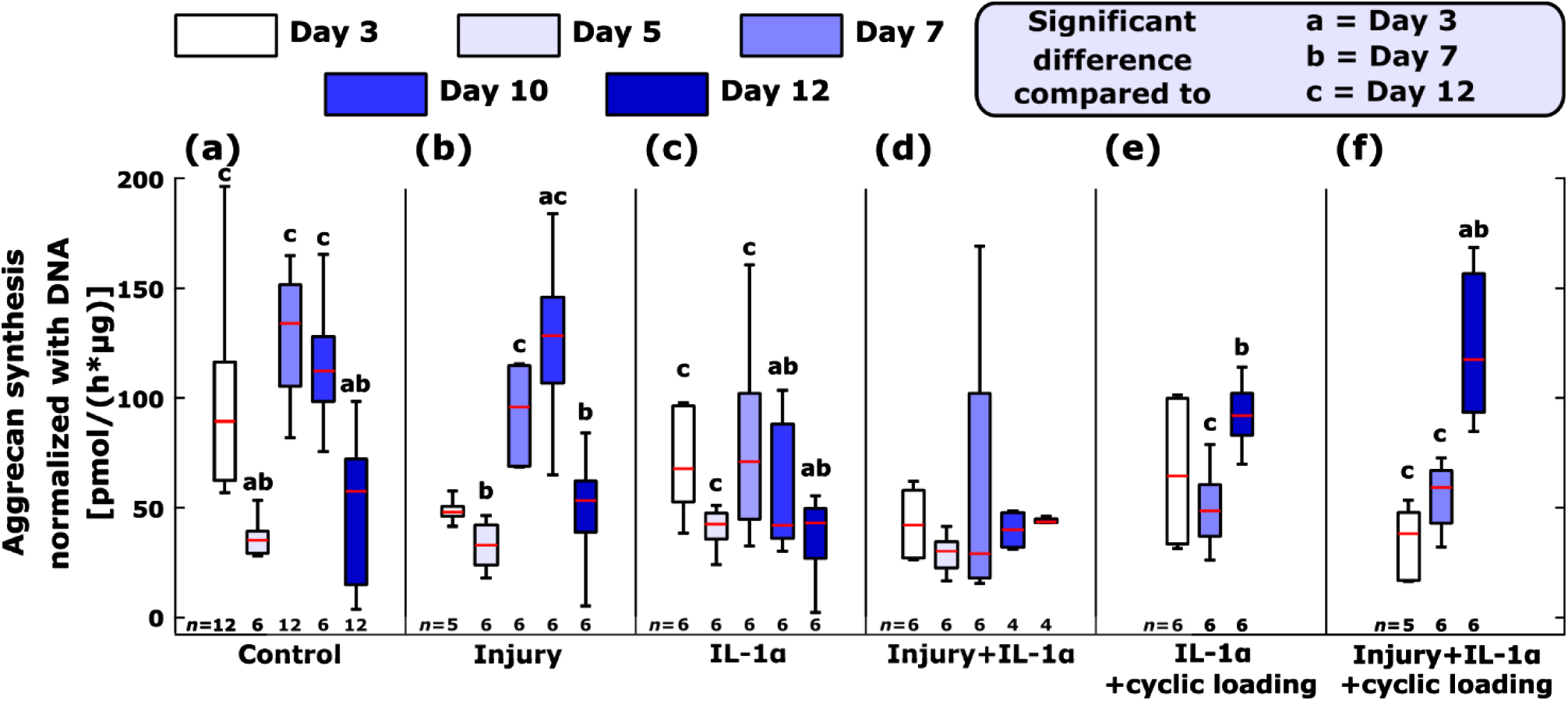
Aggrecan biosynthesis (^35^S incorporation) normalized with the amount of DNA left in the chondrocytes in different treatment groups at different time points. Biosynthesis rates were measured for **(a)** free-swelling control (CTRL), **(b)** injury-only (INJ), **(c)** interleukin-1α-challenged (IL), **(d)** injured and IL-1α-challenged (INJ-IL), **(e)** IL-1α-challenged and cyclically loaded (IL-CL), and **(f)** injured, IL-1α-challenged, and cyclically loaded group (INJ-IL-CL). IL-1α decreased aggrecan biosynthesis significantly over time. Chondrocytes in the INJ group increased aggrecan biosynthesis transiently with a 7–10-day delay from the time of injury, but this was inhibited in the INJ-IL group when exogenous IL-1α was present. In turn, cyclically loaded plugs showed late (12 days after beginning of the treatment) and expedited aggrecan biosynthesis even in the presence of IL-1α and chondral lesions. The amount of plugs *n* designated for each time point is shown on the bottom of each subpanel. The letters above the boxplots indicate statistically significant differences in aggrecan biosynthesis rate compared to that at 3, 7, and 12 days after initiation of the treatment (*p* < 0.05; linear mixed effects model).

Injurious loading and cytokines separately and simultaneously reduced aggrecan biosynthesis significantly over seven days with respect to CTRL (day 7 median biosynthesis rates CTRL 134 pmol/h/μg, INJ 96 pmol/h/μg, IL 71 pmol/h/μg, INJ-IL 29 pmol/h/μg; CTRL *vs.* INJ *p* = 0.010, CTRL *vs.* IL *p* = 0.001, CTRL *vs.* INJ-IL *p* < 0.001; S1 Supplementary Material, Subsection S2.5, Fig. S10c). The significant increase in cyclic loading-modulated aggrecan biosynthesis occurred later on day 12 (median biosynthesis rates CTRL 57 pmol/h/μg, IL-CL 92 pmol/h/μg, INJ-IL-CL 118 pmol/h/μg; CTRL *vs.* IL-CL *p* = 0.001, CTRL *vs.* INJ-IL-CL *p* < 0.001; Fig. S7e). See more details in the Supplementary Material S1 (Subsection S2.5, Fig. S10, Tables S13-S15).

### 2.5 Cell death caused by chondral injuries was localized while IL-1α caused more widely spread cell death, mitigated by cyclic loading only in the deeper cartilage

Qualitatively, we observed cell death especially close to the cartilage injuries in groups involving an injurious compression (Fig. 8). In groups involving an IL-1α-challenge, cell death was pronounced close to all edges but it also reached deeper and more central layers of the slices. We also observed similar areas of cell death throughout the plug but concentrating near the edges in some of the CTRL samples; we assume the latter was probably caused by the biopsy punch or cutting of the histological sections. In the INJ-IL group, we observed marked cell death near lesions and at the immediate regions next to the lesions as opposed to tissues further away from them (see *e.g.* the top surface areas in the INJ-IL group, day 7, Fig. 8). Some of our cyclically loaded samples (such as day 12 sample from the IL-CL group) showed qualitatively more cell death near superficial layers and more viable cells in the deeper cartilage than samples without cyclic loading (such as day 12 IL group). Also, some of our day 7 plugs from the INJ-IL-CL group showed qualitatively similar or slightly increased cell death in the damaged superficial areas but increased cell viability in the deeper regions compared to the INJ-IL group.

**Fig. 8.**
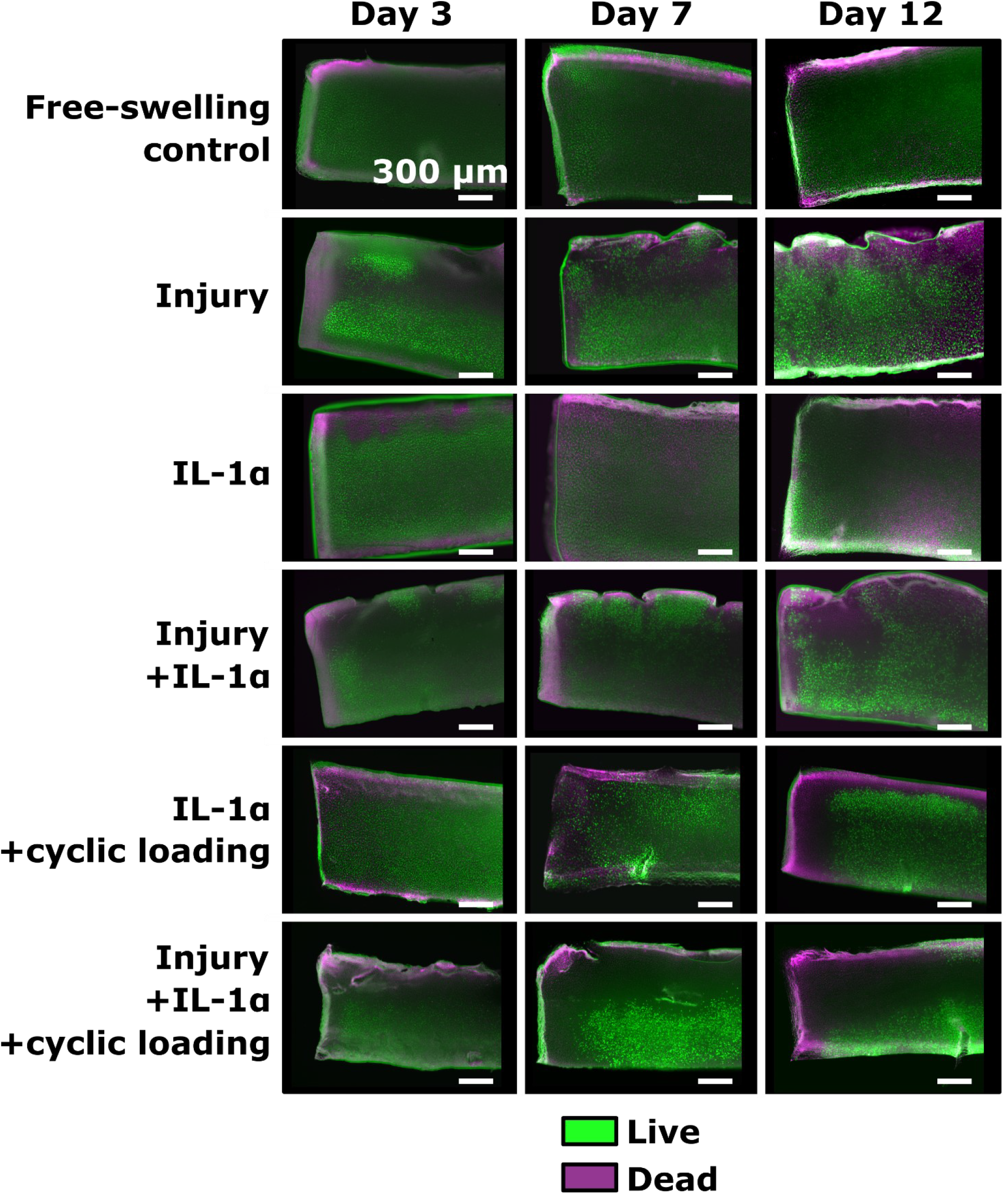
Cell viability in histological slices after different culture conditions. Green color (fluorescein diacetate stain) shows live cells and magenta color (propidium iodide stain) dead cells. Cell death was noticeable near chondral lesions. In interleukin (IL)-1α-challenged samples, cell death was pronounced near explant edges but reaching also the deeper layers of cartilage. However, cyclically loaded samples could still show vital cells on day 12 but only in the deep cartilage. Each treatment and time point combination shown here had two plugs prepared for cell viability analysis (2–3 slices per plug from the middle of the plug), except the free-swelling control group which had four plugs prepared.

## 3 Discussion

In summary, we modeled *in vitro* the combined effect of injuries, exogenous pro-inflammatory IL-1α-challenge, and moderate cyclic loading on GAG loss, aggrecan biosynthesis, and cell viability in young calf cartilage. The main findings were: **(1)** accumulated GAG loss increased dose-dependently with increasing IL-1α concentration, *i.e*., inflammatory challenge (Fig. S2). **(2)** Cyclic loading (15% strain amplitude, 1 Hz) ameliorated the (normalized) accumulated GAG loss during the first 2–4 days of treatment (Fig. 3) by protecting especially the deep cartilage regions (Fig. 5). **(3)** At the end of the 12-day experiments, the beneficial effects of the chosen cyclic loading regime had been compromised; this cyclic loading could occasionally inhibit — though not fully protect — cartilage from acceleration of accumulated GAG loss (Fig. 2). In turn, normalized accumulated GAG loss could even increase compared to samples that were otherwise similarly treated but not cyclically loaded (Fig. 3). However, cyclic loading could still slow down the rate of localized GAG depletion away from lesions (Figs. 6, S5, S6, S8, and S9). **(4)** Regarding aggrecan biosynthesis, injury caused a transient increase, IL-1α decreased the aggrecan biosynthesis, and the applied moderate cyclic loading increased biosynthesis only at 12 days of loading (Fig. 7). **(5)** Cell death was localized close to lesions in injured plugs and was notable over larger areas when cytokines were present, whereas deep cartilage of cyclically loaded plugs could retain cell viability (Fig. 8). **(6)** Near the lesions, GAG loss was localized to the same areas as cell death (Figs. 4 and 8). Taken together, for injured and IL-1-inflamed bovine cartilage, our cyclic loading scheme was **(1)** beneficial early (days 2–4) but **(2)** later-on (days 8–12) its pro-anabolic effects were severely hampered or questionable at best – the bulk GAG loss might not be decreased though biosynthesis would increase and deep cartilage GAG content and viable cells would be partially preserved. These novel findings are essential for a deeper understanding of the time-dependent and depth-dependent early stages of PTOA pathophysiology and for the development of countermeasures for PTOA progression, such as designing rehabilitation procedures, disease-modifying drugs, and high-biofidelity computational models offering decision-making guidance for treatment.

We consider our choice of 1 ng/ml of IL-1α as a relatively high cytokine concentration which mimics the slow PTOA progression rapidly in an experimental setting. For comparison, physiological/mildly inflamed human synovial fluid has 0.01–0.25 ng/ml of IL-1α^36, 37^ and 0.005–0.02 ng/ml of isoform IL-1β^36, 38^.

Our findings are generally in line with the current literature. GAG loss has previously been found to be cytokine concentration-dependent^10, 18, 19, 39^, which was confirmed by our findings. Furthermore, chondral injuries aggravate cytokine-modulated degradation^5, 10, 35^, for which our results give further support. We noted a sharp increase in accumulated GAG loss after injurious loading, also observed elsewhere^6–9, 22^. Localized GAG loss was concentrated near the lesions similar to a previous study^11^. Surprisingly, as opposed to pro-catabolic cytokine challenge of IL-6 and TNFα^5^, our data demonstrate that although moderate cyclic loading (such as 10 and 20% strain amplitudes applied with 0.5 Hz frequency, 1h on/5h off per day in the work of Li *et al.*^5^) is widely considered to be beneficial for articular cartilage health^5, 32–34, 40, 41^, the cyclic loading of our protocol could result in an increase (Fig. 3bcde) or only modest decrease (Fig. 2) in the 8–12-day accumulated GAG loss caused by the combination of injurious loading and relatively high concentration of IL-1α. However, we acknowledge that other cyclic loading protocols (like decreasing the strain amplitude to 10%) could lead to different results and might decrease GAG loss even after 12 days and not just in the early days. However, the inability of cyclic loading to counteract the long-term deleterious effects of IL-1 was also observed in an experiment with chondrocyte-seeded agarose constructs^42^. In turn, when considering the early 1–3-day response, previous studies^32–34^ concluded that the degradative effects of IL-1 could be abolished with cyclic loading in terms of GAG loss, supported by our findings (Fig. 3de).

Pro-catabolic cytokines effectively inhibit aggrecan biosynthesis and endogenous repair mechanisms^5, 43^ (Fig. 7c). The experiments of Li *et al.*^5^ suggest that injuries could increase biosynthesis in inflamed cartilage, but our data hints that combination of injury and IL-1 decreases biosynthesis synergistically or the presence IL-1 thwarts the injury-related efforts to increase biosynthesis (S1 Supplementary Material, Subsection S2.5, Fig. S10d). We noted a transient increase in aggrecan biosynthesis in the INJ group (Fig. 7b). Similar increase of ^35^S-sulfate incorporation during early disease stages and later followed by a decrease of sulfate incorporation has been observed in damaged human articular cartilage as well^44^. Thus, an injury might induce the viable chondrocytes close to lesions and in the deeper zones to attempt an early compensation for the GAG loss. However, the GAG content on newly synthesized aggrecan could be abnormal with increased ratio of chondroitin-4-sulfate to total chondroitin sulfate content^12, 13^. We also provide further supporting evidence that moderate cyclic loading increases aggrecan production^5, 45^ (Fig. 7ef). Our data also shows that the beneficial effects of the proposed cyclic loading scheme in terms of increased aggrecan biosynthesis are not attained instantaneously, but after a 12-day delay from the beginning of treatment (Fig. 7ef). Aggrecan biosynthesis also decreases with chondrocyte death; apoptosis has been previously observed to occur near chondral lesions^5, 11^ as well as to be pronounced towards the inner parts of inflamed cartilage^5^, like our biosynthesis findings (Fig. 7d) and obtained histological sections also demonstrate (such as Fig. 8, day 12, INJ-IL group). However, chondrocytes in cyclically loaded plugs could still be viable and synthesize new matrix constituents in the deeper regions^27, 46^ (Fig. 8, day 7, INJ-IL-CL group).

The reasons behind the obtained GAG loss results are manifold and not fully understood. A previous study suggested that over short time periods (24 h after injury), the injury-related GAG loss occurs due to microdamage rather than pro-catabolic proteases synthesized by nearby chondrocytes^47^. We speculate that the loss of structural integrity^7^ (caused by injurious loading and high shear strains near lesions^48^) might facilitate diffusion of GAG fragments out of cartilage and cytokine diffusion into cartilage through lesion surfaces (larger surface area to diffuse through compared to intact cartilage). Enhanced cytokine transport could cause further GAG depletion^10, 35^ via increased aggrecanase expression^20, 21^ near lesions. Moreover, during or after cyclic loading, the influx of fluid back into cartilage might also facilitate advection of cytokines into tissue^49^ especially in the superficial and injured areas; similar increase in cytokine infiltration has been observed in cyclically loaded and TNFα-cultured intervertebral discs^50^. Cyclic loading-induced elevation of fluid velocities might also promote depletion of resident or newly synthesized proteoglycans from the tissue^11^. Lastly, GAG loss is also associated with decrease of aggrecan biosynthesis due to cell death. Besides causing cell death themselves^5, 14^, injuries and cytokines could promote oxidative stress^15^ and expression of pro-catabolic nitric oxide^51^ which cause further cell death^16, 17^.

We found that moderate cyclic loading could be beneficial initially but might not prevent degradation of injured and IL-1-inflamed cartilage over extended periods of time; instead, cyclic loading of a specific type could even become detrimental especially if relatively high concentrations of IL-1-cytokines are present. Our results provide a new hypothetical intervention for PTOA progression and a future topic for *in vivo* investigation: patients with an injured or surgically operated knee joint would be medicated early-on with drugs counteracting pro-catabolic cytokines as well as undergo physical rehabilitation as soon as possible. However, the rehabilitation period should not take place when pro-inflammatory cytokine levels remain chronically elevated. Despite the unquestionable benefits of physical rehabilitation in terms of reduced pain and increased joint functionality^52^, randomized controlled clinical trials also report that the physical exercise itself (*e.g.*, not combined with dietary intervention) does not have an effect on the levels of inflammatory biomarkers or decreases them only moderately^53, 54^. Therefore, while rehabilitation might not be able to restrain inflammation, the cyclic loading of the joint during physical activities could pose a risk of enhancing cytokine transport into cartilage via advection^49^, possibly contributing to PTOA progression. This could be one of the mechanisms why inhibiting chronic inflammation is crucial. Using steroidal or non-steroidal anti-inflammatory drugs (NSAID) would appear attractive based on our results, but there is widespread evidence of their potentially deleterious effects on cartilage especially with high doses and long-term usage^55, 56^. Thus, novel therapeutic interventions would be necessary to both avoid the well-known risks of NSAID and to counteract the pro-catabolic cytokine responses. Specifically, IL-1 receptor antagonists^57, 58^, cytokine synthesis inhibitors^59, 60^ and human blood-based solutions^61^ could work as a means to reduce IL-1-mediated disease progression. Though some drugs may clinically mitigate symptoms in rheumatoid arthritis patients but not in PTOA patients^58, 62^, this does not conclusively rule out their potential to decelerate cartilage degeneration especially when delivered in tandem with well-timed rehabilitation.

Our main study limitation is the relatively small sample size (*n* = 6 analyzed plugs per treatment group and time point combination), especially at later time points. Furthermore, we did not study collagen loss since earlier animal studies^63^ propose that GAG loss precedes collagen loss and notable collagen depletion is exhibited only after 12 days of culturing with IL-1α^18^. Additionally, we did not include a group for **(1)** cyclic loading-only and **(2)** combined injurious and cyclic loading, since those have already been studied although without reporting the accumulated GAG loss and changes in aggrecan biosynthesis^11^.

In the future, models of experimentally induced PTOA as used here are of utmost importance to gain knowledge of the earliest disease stages and for the development of methods tackling the disease. In addition to IL-1, TNFα and IL-6 are believed to be major players in PTOA progression for example after anterior cruciate ligament rupture^36, 64^; therefore, one interesting topic to study is to use these cytokines simultaneously with upscaled concentrations but in proportions as found *in vivo*. The cyclic loading and culture could possibly be continued for a longer period than 12 days after injury to confirm the longer-term effects of cyclic loading. The cyclic loading protocol itself should also be scrutinized to find how different loading parameters (strain amplitude, loading frequency, on/off timing, exerting controlled shear strains as well rather than single-axis compression) might change the short- and long-term cartilage response. Also, further investigation of the mechanisms of how and why prolonged loading becomes detrimental after the initial benefit is of particular interest; answers to these questions could be helpful for designing enhanced osteoarthritis therapies. Moreover, on this experimental platform, the inflammation could be mediated by a host of different anti-inflammatory or disease-modifying drugs.

## 4 Methods

### 4.1 Bovine cartilage harvest and culture conditions

Bovine cartilage plugs were harvested from the patellofemoral grooves (femoral surface) of freshly slaughtered 1–2-week-old calves (Research 87 Inc., Boylston, MA) similarly to previous studies^11, 18, 65^. A total of six knee joints from six animals were used for the main experiments and additional two joints from two different animals were utilized for the pilot study. Cylindrical cartilage plugs were obtained with a 3-mm-diameter dermal punch and washed with Dulbecco’s phosphate buffered saline (D-PBS) supplemented with 1% penicillin G-streptomycin-amphotericin B (PSA; 100 units/ml, 100 μg/ml, 0.25 μg/ml, respectively) without calcium and magnesium. Subsequently, the plugs were trimmed with a razor blade to obtain 1-mm thick cartilage plugs including the intact superficial layer.

Cartilage plugs were equilibrated for two days before the experiments (37 **°**C, 5% CO_2_; Fig. 1a) submerged in serum-free culture medium consisting of low-glucose (1 g/l) Dulbecco’s Modified Eagle’s medium (DMEM; Sigma-Aldrich, MilliporeSigma, Burlington, MA) supplemented with 1% PSA, 1% insulin-transferrin-selenium (ITS), 10 mM 4-(2-hydroxyethyl)-1-piperazineethanesulfonic acid (HEPES) buffer, 0.1 mA non-essential amino acids (NEAA), 0.4 mM proline, and 20 μg/ml ascorbic acid. Following the equilibration period, the medium was changed every two days during the experiments.

### 4.2 Experimental groups and time points

Cartilage plugs were subjected to different treatment conditions for up to 12 days after the equilibration period (Fig. 1a). The experimental groups and the treatment conditions were **(1)** free-swelling control (CTRL), **(2)** injury-only (INJ), **(3)** IL-1α-challenge-only (IL), **(4)** injured and IL-1α-challenged (INJ-IL), **(5)** IL-1α-challenged and cyclically loaded (IL-CL), and **(6)** injured, IL-1α-challenged, and cyclically loaded (INJ-IL-CL). In total, we harvested 406 cartilage plugs from six different bovine knees. The CTRL group was assigned 112 plugs (48 to GAG loss and biosynthesis analyses, 48 to digital densitometry, and 16 to cell viability); INJ, IL, and INJ-IL groups were each assigned 70 plugs (30 to GAG loss and biosynthesis analyses, 30 to digital densitometry, and 10 to cell viability); IL-CL and INJ-IL-CL groups were both assigned 42 plugs (18 to GAG loss and biosynthesis analyses, 18 to digital densitometry, and 6 to cell viability). These plugs were then assigned to different time points.

The culture was terminated on days 3, 5, 7, 10, or 12 after initiation of the treatment for the CTRL group and the groups without cyclic loading (INJ, IL, INJ-IL), and on days 3, 7, and 12 for the CTRL group and the cyclically loaded groups (IL-CL, INJ-IL-CL; Fig. 1b). The CTRL samples were pooled. In total, samples from four animals were assigned to the groups without cyclic loading and samples from two different animals to the cyclically loaded groups (Table 1). Ultimately, 14 plugs were assigned for each treatment group and time point combination (6 to GAG loss and biosynthesis analyses, 6 to digital densitometry, and 2 to cell viability). Location-matching of the plugs was done by assigning samples from all four areas along the patellofemoral surface to each treatment group. The areas were **(1)** medio-distal, **(2)** latero-distal, **(3)** medio-proximal, and **(4)** latero-proximal; the division of the joint surface to the medio-lateral and proximo-distal parts was approximated from the middle point of the respective directions (Fig. 1a). For (normalized) accumulated GAG loss and aggrecan biosynthesis assays, two plugs were assigned to all treatment groups from area 1, one plug from area 2, two plugs from area 3, and one plug from area 4. For localized GAG loss analyses (optical density profiles and regional averages), one plug was assigned to all treatment groups from area 1, two plugs from area 2, one plug from area 3, and two plugs from area 4. Taken together, for all GAG analyses, each group had three plugs from all four areas. For qualitative assessment of cell viability, the plugs were taken from the areas where cartilage was still available for harvesting.

**Table 1.** Assigning cartilage plugs from different animals to the treatment groups without cyclic loading and cyclically loaded groups. Treatment groups without cyclic loading (injury-only; IL-1α-challenge-only; injured and IL-1α-challenged) were assigned four animals which provided samples for five time points; 3, 5, 7, 10, and 12 days after initiation of the treatment. In experiments with cyclically loaded groups (IL-1α-challenged and cyclically loaded; injured, IL-1α-challenged, and cyclically loaded), we used two animals providing cartilage plugs for three time points; days 3, 7, and 12 after initiation of the treatment. Each animal provided also free-swelling control plugs. Furthermore, the glycosaminoglycan content lost to culture medium was measured at several time points; for instance, plugs whose culture was terminated on day 10 provided data for days 0, 2, 4, 6, 8, and 10 (culture medium was changed every two days).

| Animal | Cyclic loading | Culture termination days |
| --- | --- | --- |
| 1 | No | 10, 12 |
| 2 | No | 7, 10, 12 |
| 3 | No | 3, 5, 7 |
| 4 | No | 3, 5 |
| 5 | Yes | 3, 7, 12 |
| 6 | Yes | 3, 7, 12 |

### Injurious loading

On day 0, *i.e.* after the 2-day equilibration period, the cartilage plugs in groups involving injurious loading were subjected to a single load–unload cycle of radially unconfined compression with 50% axial strain with respect to the sample-specific thickness (strain rate 100%/s) within a custom-designed incubator-housed loading apparatus^66^. The resulting peak stresses were in the range of 15–25 MPa. This loading scheme is known to produce cracks in articular cartilage samples^5, 11^.

#### Cytokine culture

Cytokine-challenged plugs were cultured in the same culture medium as described in Subsection 4.1 but supplemented with exogenous pro-inflammatory cytokine interleukin (IL)-1α, thus allowing diffusion of cytokines into the cartilage. Based on the pilot study involving different concentrations of IL-1α, a concentration of 1 ng/ml was chosen for cytokine-cultured treatment groups (see Subsection 2.1 and S1 Supplementary Material, Subsections S2.1-S2.2, Figs. S1 and S2). This concentration has also been used previously^18, 32^.

#### Cyclic loading

Cyclically loaded plugs were placed in a 30-well polysulfone loading chamber (the plugs that did not have room in the 30-well chamber were placed in an additional 12-well chamber) and subjected to 10% compressive offset strain (achieved with five 2% axial strain ramp-and-hold increments, each increment followed by a six-minute stress-relaxation period) to ensure proper, repeatable, and similar contact between all the samples and the chamber. Subsequently, the plugs were cyclically loaded in unconfined compression with 15% axial strain (1 Hz, haversine waveform) continuously for 1h with an incubator-housed cyclic loading apparatus, followed by a resting period in another incubator (the chamber lid was slightly elevated to avoid static loading of cartilage plugs due to lid weight)^5, 11^. The cyclic loading–resting-cycle was repeated four times a day with one cyclic loading device, resting times being 3h, 4h, 3h, and 10h (overnight rest) between loadings within one 24-hour period, thus mimicking normal everyday activities (see Supplementary Material S1, Section S1, Fig. S1).

### 4.3 Assays

Four biochemical assays and digital densitometry were utilized to quantify the effects of injurious loading, pro-inflammatory cytokine culture and cyclic loading on articular cartilage glycosaminoglycan (GAG) content, DNA content, aggrecan biosynthesis rate, and chondrocyte viability (Fig. 1c).

#### Accumulated glycosaminoglycan loss

Dimethylmethylene blue (DMMB) dye binding assay^67^ was used to quantify the accumulative loss of sulfated GAG from the whole plugs. GAG loss was measured from the culture medium (that was changed every two days) and, after digestion with proteinase K over 24 h, from the plugs at termination days of the experiments. The accumulated GAG loss results are presented over time relative to the initial total GAG content in the plugs (*i.e.*, GAG depleted into the culture medium and retained in the plug):

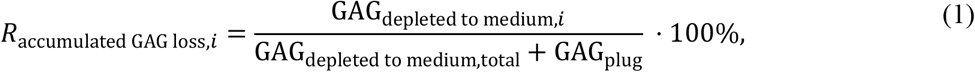

where *R*_accumulated GAG loss,*i*_ is the accumulated GAG loss on day *i* relative to the initial total GAG content, GAG_depleted to medium,*i*_ is the accumulated GAG content (μg) depleted to the medium from the beginning of the culture until day *i*, GAG_depleted to medium,total_ is the total amount of GAG (μg) depleted to the medium during the whole experiment, and GAG_plug_ is the amount of GAG left inside the plug (μg).

#### Normalized accumulated glycosaminoglycan loss

Accumulated GAG loss (Eq. 1) was calculated in all the groups. Next, this data from all the treated plugs was normalized with that of animal- and location-matched CTRL plugs in order to obtain a fold change measure of proteoglycan matrix degradation. The normalization was utilized in order to detect the GAG loss changes in response to different treatments without a bias from animal or cartilage location-related differences in GAG depletion. Furthermore, we noted that the GAG release differed between animals, thus underlining the necessity of such normalization; the average accumulated GAG loss of controls of groups without cyclic loading was significantly higher than that of controls of cyclically loaded groups (*p* < 0.05) on days 8, 10 and 12. This suggests that the non-normalized accumulated GAG loss would be overestimated for instance for the INJ-IL group compared to the INJ-IL-CL group. The normalized accumulated GAG loss was calculated as a fold change between the accumulated GAG loss (%, Eq. 1) of the treated plug and that of the corresponding control plug:

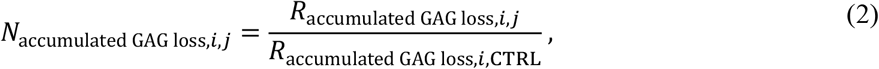

where *N*_accumulated GAG loss,*i*,*j*_ is the normalized accumulated GAG loss on day *i* for a plug in group *j* (*j* = INJ, IL, INJ-IL, IL-CL, or INJ-IL-CL), *R*_accumulated GAG loss,*i*,*j*_ is the accumulated GAG loss (%) on day *i* for a plug in group *j*, and *R*_accumulated GAG loss,*i*,CTRL_ is the accumulated GAG loss (%) on day *i* for the corresponding (same animal and location) control plug.

#### Localized glycosaminoglycan loss

On the termination day of the experiments, the samples designated to localized GAG loss analysis were fixed with neutral buffered 10% formalin solution. Next, the plugs were dehydrated with graded series of alcohols, transferred to intermediary xylene, and finally embedded in paraffin. Nine histological sections (thickness 3 μm) were cut perpendicular to the surface of the plugs and stained with Safranin-O. Ultimately, three representative slices^68^ per plug were chosen (*e.g.*, badly ruptured slices were discarded) for analysis of optical density, a proxy measure of GAG content^69, 70^. Optical density was quantified with digital densitometry^68^ by using a conventional light microscope (Nikon Microphot FXA, Nikon Inc., Tokyo, Japan), monochromator (wavelength *λ* = 492 ± 5 nm), and a charge-coupled device camera (Hamamatsu ORCA-ER, Hamamatsu Photonics, Hamamatsu, Japan). The obtained grayscale images (4x magnification, pixel size 1.23 μm) were converted to optical density maps by using calibration images of filters with known optical densities of 0.0, 0.15, 0.3, 0.6, 1.0, 1.3, 1.6, 2.0, 2.3, 2.6, and 3.0 (Schott, Mainz, Germany).

##### Optical density profiles

Two depth-wise optical density profiles were obtained from each slice (*i.e.*, six profiles per plug in total). For injured samples, 200 μm wide profiles were taken away from injuries from both sides of them; this width was chosen to ensure that the profiles could be taken from each slice without overlap with lesions. For uninjured samples, the profiles were taken **(1)** from the middle and **(2)** between the middle and lateral edge of each slice. Each profile was averaged perpendicular to the depth-wise direction. Then, the profiles on each three slices per plug were averaged, ultimately leading to two average profiles per plug.

##### Average regional optical densities

Regions away from injuries (width 200 μm, depth 150 μm) were chosen from the superficial zone at the same locations as the optical density profiles, *i.e.*, two away-from-injury regions per slice. Furthermore, one region within 50 μm from a lesion edge was obtained by selecting lesion edges and using morphological dilation in MATLAB (R2017b, Image Processing Toolbox, The MathWorks Inc., Natick, MA, USA). These distances were chosen to restrict our analysis to the superficial/transitional zone, where the most rapid GAG depletion would most likely occur, and during preliminary tests a distance of 50 μm around lesions would approximately reach the same 150 μm depth as the regions away from lesions in majority of the cartilage slices. Average optical density was calculated within these regions per slice. Finally, by averaging the optical densities of corresponding regions on the three slices we obtained sample-specific estimations of localized GAG content.

##### DNA assay

The DNA content of the plugs was assayed with fluorometric Hoechst 33258 dye binding assay^71^ after digestion with proteinase K. The results of DNA assay are not presented explicitly, instead they were used to normalize the aggrecan biosynthesis rates in order to obtain quantitative biosynthesis data with respect to actual amount of DNA in the plugs instead of plug wet weights (which is affected by the drying method of the plug, ECM loss and biosynthesis).

##### Aggrecan biosynthesis

The aggrecan biosynthesis rates were quantified with ^35^S-sulfate incorporation assay after digestion with proteinase K. Cartilage plugs were cultured in medium supplemented with 5 μCi/ml of ^35^S-sulfate radiolabel for the last 48 hours of experiments (24h for day 3 plugs) and upon termination washed with D-PBS for 4x15min. The duration of exposure to radiolabel was used to normalize the rate of radioactive sulfate incorporation into sulfated GAG^18, 65^. The aggrecan biosynthesis results were normalized with sample-specific DNA content.

##### Cell viability

Distribution of live and dead cells was assessed visually from stained histological sections. 100–300 μm thick slices were cut from the middle of the plugs perpendicular to the top surface of the plugs with a scalpel. Then, the slices were incubated in cell viability assay solution for 3 min covered from light sources (1 ml D-PBS, 2 μl fluorescein diacetate [FDA; staining viable cells green], and 4.6 μl propidium iodide [PI; staining non-viable cells red and transformed into magenta with ImageJ]). Subsequently, the slices were washed thrice with PBS (3 x 3 min). The slices were imaged with Nikon fluorescence microscope through two filters (to detect fluorescence from FDA and PI separately) at 4x magnification^11, 18^.

#### 4.4 Statistical analyses

All statistical analyses were conducted with linear mixed effects (LME) model for (normalized) accumulated GAG loss, localized GAG loss, and aggrecan biosynthesis in IBM SPSS Statistics 25.0 (SPSS Inc., IBM Company, Armonk, NY, USA). Data points more than 1.5 times the interquartile range below the 25% quartile or above the 75% quartile were discarded as outliers. The LME model considered different animals as subjects with random intercepts, and treatment, time, and areas on the patellofemoral surface as fixed factors. Accumulated GAG loss assay provided repeated measures data at time points when culture medium was collected; we assumed that the model error terms within plugs were correlated over time. Thus, we chose first-order autoregressive covariance (AR(1)) as a repeated covariance type. We divided the optical density profiles to 100 depth-wise points to compare the GAG content of different treatment groups at different normalized depths. This depth-wise comparison was done conservatively by observing whether the predicted mean of the modeled variable falls into the 95% confidence interval of the variable under comparison (non-significant difference) or not (significant difference). With the analysis of optical density profiles and regional averages, the location-dependent variation within samples (*i.e.*, two profiles/regions away from lesions) was considered in the LME model. Localized GAG content and aggrecan biosynthesis data analyses were done similarly as (normalized) accumulated GAG loss assay but without repeated measures design. The significance level for pair-wise comparisons was set at *α* = 0.05, and the post-hoc analyses were made with Fisher’s Least Significant Difference.

## Supporting information

**S1 Supplementary Material.** Electronic supplementary material containing additional and more detailed information about the underlying experimental findings, including statistics.

**Fig. S1.** The cyclic loading protocol mimicking daily activities.

**Fig. S2. Accumulated glycosaminoglycan (GAG) loss in the pilot study with different concentrations of interleukin (IL)-1α to find a suitable concentration for the rest of the study.**

**Fig. S3. Aggrecan biosynthesis in the pilot study with different concentrations of interleukin (IL)-1α to find a suitable concentration for the rest of the study.**

**Fig. S4. Normalized accumulated glycosaminoglycan (GAG) loss between different treatment groups at different time points.**

**Fig. S5. Optical density profiles and comparison between groups on days 3 and 7 in more detail.**

**Fig. S6. Optical density profiles on days 0, 5, 10, and 12.**

**Fig. S7. Further comparison of optical density profiles on day 3.**

**Fig. S8. Localized glycosaminoglycan (GAG) content near and away from lesions on days 3 and 7.**

**Fig. S9. Localized glycosaminoglycan (GAG) content near and away from lesions on days 0, 5, 10, and 12.**

**Fig. S10. Aggrecan biosynthesis (^35^S incorporation) normalized with the amount of DNA left in the chondrocytes between different treatment groups at different time points.**

**Table S1. Accumulated glycosaminoglycan (GAG) loss in the pilot study.**

**Table S2. Statistics of accumulated GAG loss between different time points within groups in the pilot study.**

**Table S3. Statistics of accumulated GAG loss between different groups at several time points in the pilot study.**

**Table S4. Aggrecan biosynthesis rate in the pilot study.**

**Table S5. Statistics of aggrecan biosynthesis rate between different time points within groups in the pilot study.**

**Table S6. Statistics of aggrecan biosynthesis rate between different groups at several time points in the pilot study.**

**Table S7. Accumulated glycosaminoglycan (GAG) loss.**

**Table S8. Statistics of accumulated GAG loss between different time points within groups.**

**Table S9. Statistics of accumulated GAG loss between different groups at several time points.**

**Table S10. Normalized accumulated glycosaminoglycan (GAG) loss.**

**Table S11. Statistics of normalized accumulated GAG loss between different time points within groups.**

**Table S12. Statistics of normalized accumulated GAG loss between different groups at several time points.**

**Table S13. Aggrecan biosynthesis rate.**

**Table S14. Statistics of aggrecan biosynthesis rate between different time points within groups.**

**Table S15. Statistics of aggrecan biosynthesis rate between different groups at several time points.**

## Acknowledgements

The authors appreciate the support of the University of Eastern Finland and the Massachusetts Institute of Technology to conduct this study. Yamini Krishnan, Ph.D., is acknowledged for help in the laboratory and guidance during pre-processing of the raw data. Eija Rahunen (Institute of Biomedicine, Cell and Tissue Imaging Unit, UEF) is acknowledged for the processing of the samples designated for digital densitometry analysis. Kalle Karjalainen, M.Sc., and Mohammedhossein Ebrahimi, M.Sc., are acknowledged for the technical support for the digital densitometry analysis.

## Author contributions

ASAE contributed to the design of the study and experiments, carrying out the experiments, data analysis and interpretation of data, and drafting and revising the manuscript. CF contributed to the design of the study and experiments, interpreting the data, drafting and critical revision of the article for intellectual content. PT contributed to the design of the study and experiments, data analysis and interpretation of data, drafting and critical revision of the article for intellectual content. HKH contributed to the design of the experiments, interpreting the data and critical revision of the article for intellectual content. EHF contributed to the design of the experiments, interpreting the data and critical revision of the article for intellectual content. SM contributed to interpreting the data, statistical analysis and critical revision of the article for intellectual content. PN contributed to interpreting the data and critical revision of the article for intellectual content. PJ contributed to the design of the study, interpreting data, drafting and critically reviewed the article for important intellectual content. AJG contributed to the conception and design of the study, experiments, data analysis, interpretation of data, drafting and critical revision of the article for intellectual content. RKK contributed to the conception and design of the study, experiments, data analysis and interpretation of data, drafting and critical revision of the article for important intellectual content.

## Data availability

The datasets generated and analyzed during this study are available from the corresponding author on a reasonable request.

