## Supplementary material for "Cyclic loading regime considered beneficial does not protect injured and interleukin-1-inflamed cartilage from post-traumatic osteoarthritis": S1 Supplementary Material

### Electronic Supplementary Material

### S1 Methods

The Fig. S1 provides a schematic of the used cyclic loading protocol.

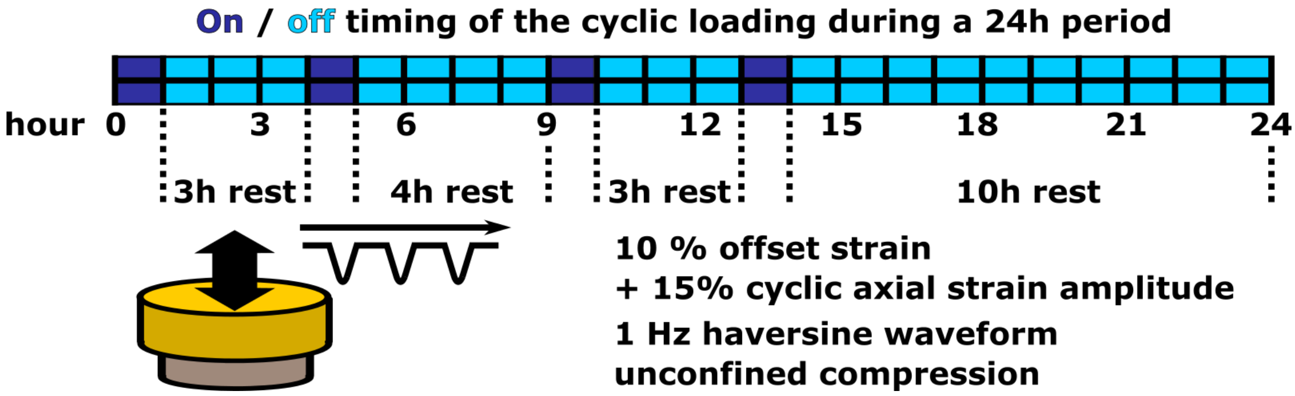


**Fig. S1. The cyclic loading protocol mimicking daily activities.** During a 24h period, the cartilage plugs were subjected to compressive cyclic loading in unconfined compression for four 1h periods, separated by 3h, 4h, 3h, and 10h (overnight) rest periods. A loading period commenced with a 10% offset strain (five 2% axial strain ramp-and-hold increments with 6-min stress-relaxations between increments) to ensure proper, repeatable, and similar contact between all the samples and the chamber. Next, the samples were cyclically loaded with 15% axial strain amplitude (1 Hz, haversine waveform) continuously for 1h with an incubator-housed cyclic loading apparatus. During a rest period, the samples were in another incubator without mechanical loading.

### S2 Results

All statistical analyses were carried out with linear mixed effects models. The supplementary tables S1-S15 show the estimated marginal means and statistical comparisons based on those.

#### S2.1 High exogenous cytokine concentration leads to high proteoglycan matrix damage

In the pilot study, 10 ng/ml of IL-1α caused significantly more accumulated GAG loss compared to the other groups (free-swelling control (CTRL) and IL-1 concentrations of 0.1 ng/ml, 0.5 ng/ml, and 1 ng/ml; Fig. S2, Table S1). Over time, all the groups exhibited significant GAG loss. For example, between 8 and 12 days of culture the accumulated GAG loss increased significantly in the groups with 1 ng/ml (average 15.2% on day 8 *vs.* 21.7% on day 12, $p < 0.001$) and 10 ng/ml of IL-1 (25.9% on day 8 *vs.* 50.6% on day 12, $p < 0.001$), demonstrating that the cytokine-driven degradation accelerated proteoglycan matrix damage after approximately one week of culture (Table S2). The group with 10 ng/ml of IL-1 showed higher accumulated GAG loss on days 2, 4, 6, 8, 10, and 12 compared to the group with 1 ng/ml (Fig. S2, Table S3). The concentration of 1 ng/ml depleted significantly more GAG from the plugs compared to the CTRL plugs from day 4 onwards (Fig. S2, Table S3). In Tables S1-S3, the IL-1α-challenge-only groups with concentrations of 0.1 ng/ml, 0.5 ng/ml, 1 ng/ml, and 10 ng/ml are marked as C01, C05, C1, and C10, respectively.

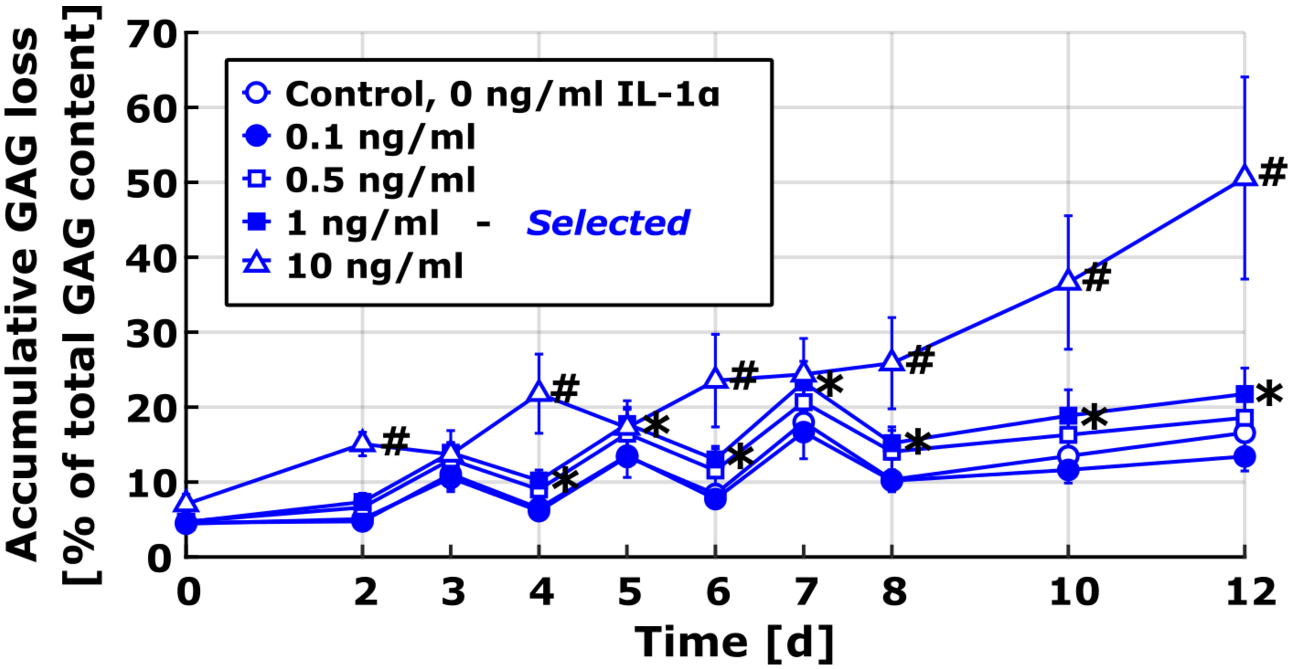


**Fig. S2. Accumulated glycosaminoglycan (GAG) loss in the pilot study with different concentrations of interleukin (IL)-1α to find a suitable concentration for the rest of the study.** Concentration of 10 ng/ml of IL-1α led to significantly greater accumulated GAG loss compared to the smaller tested concentrations (data shown as mean ± 95% confidence intervals). The IL-1α concentration of 1 ng/ml resulted in significantly higher accumulated GAG loss compared to free-swelling controls (CTRL) on days 4–12. We chose the concentration of 1 ng/ml for the rest of the study since that concentration would presumably lead to notable biochemical responses but not dominate over biomechanical factors (injury, cyclic loading). The data without location-matching of the samples was collected at different time points (days 7 and 12) from two animals (one animal for each time point, $n = 6$ plugs per group), leading to apparent decrease of average accumulated GAG loss between time points 3–4, 5–6, and 7–8 days; however, individual plugs exhibited strictly increasing GAG loss over time. Linear mixed effects model, $p < 0.05$: * = 1 ng/ml *vs.* CTRL, # = 10 ng/ml *vs.* 1 ng/ml.

**Table S1. Accumulated glycosaminoglycan (GAG) loss in the pilot study.** The results are shown as the estimated marginal mean values of accumulated GAG loss (%) with standard error from the linear mixed effect models. $N$ is the number of animals and $n$ the total amount of plugs combined from those animals. The total amount of animals in the pilot study was 2.

| Group | Day 0 $N = 2$ | 2 $N = 1$ | 3 $N = 1$ | 4 $N = 1$ | 5 $N = 1$ | 6 $N = 1$ | 7 $N = 1$ | 8 $N = 1$ | 10 $N = 1$ | 12 $N = 1$ |
|---|---|---|---|---|---|---|---|---|---|---|
| CTRL | 4.44 (1.10) $n = 12$ | 5.77 (1.18) $n = 4$ | 9.78 (1.18) $n = 6$ | 6.90 (1.28) $n = 4$ | 12.64 (1.24) $n = 6$ | 8.86 (1.32) $n = 5$ | 17.29 (1.29) $n = 6$ | 11.05 (1.39) $n = 4$ | 13.92 (1.40) $n = 5$ | 17.01 (1.43) $n = 5$ |
| C01 | 4.58 (1.10) $n = 12$ | 6.05 (1.15) $n = 5$ | 10.05 (1.18) $n = 6$ | 7.49 (1.21) $n = 6$ | 12.57 (1.24) $n = 6$ | 9.04 (1.28) $n = 5$ | 15.77 (1.29) $n = 6$ | 11.04 (1.31) $n = 6$ | 12.88 (1.34) $n = 5$ | 14.63 (1.37) $n = 5$ |
| C05 | 4.72 (1.10) $n = 12$ | 7.57 (1.14) $n = 6$ | 12.07 (1.18) $n = 6$ | 9.91 (1.21) $n = 6$ | 15.58 (1.24) $n = 6$ | 12.40 (1.26) $n = 6$ | 19.82 (1.29) $n = 6$ | 14.86 (1.31) $n = 6$ | 17.09 (1.33) $n = 6$ | 19.34 (1.34) $n = 6$ |
| C1 | 4.64 (1.10) $n = 12$ | 8.55 (1.14) $n = 6$ | 12.75 (1.18) $n = 6$ | 11.33 (1.21) $n = 6$ | 16.65 (1.24) $n = 6$ | 14.07 (1.26) $n = 6$ | 22.29 (1.29) $n = 6$ | 16.18 (1.31) $n = 6$ | 19.81 (1.33) $n = 6$ | 22.68 (1.34) $n = 6$ |
| C10 | 7.09 (1.14) $n = 11$ | 13.65 (1.21) $n = 4$ | 14.70 (1.24) $n = 5$ | 21.06 (1.24) $n = 6$ | 18.15 (1.31) $n = 5$ | 25.07 (1.31) $n = 5$ | 25.14 (1.37) $n = 5$ | 28.40 (1.35) $n = 5$ | 39.68 (1.36) $n = 5$ | 50.11 (1.36) $n = 6$ |

**Table S2. Descriptive statistics of the differences in estimated marginal means of accumulated GAG loss between different time points within groups in the pilot study.** The accumulated GAG loss is shown in the Table S1. The comparisons are shown between days 0, 2, 3, 4, 8, and 12 with 95% confidence intervals and the associated $p$-value. The $p$-values smaller than 0.05 and 0.01

are shown with light yellow and light red, respectively. The difference is shown as the mean difference in the accumulated GAG loss in the specified group in row minus the group in column.

**CTRL**

| Day | 0 | 2 | 3 | 4 | 8 |
|---|---|---|---|---|---|
| **2** | 1.33<br>(0.31; 2.35)<br>$p = .011$ | | | | |
| **3** | 5.34<br>(4.16; 6.52)<br>$p < .001$ | 4.02<br>(2.47; 5.56)<br>$p < .001$ | | | |
| **4** | 2.47<br>(0.83; 4.10)<br>$p = .003$ | 1.14<br>(-0.25; 2.53)<br>$p = .108$ | -2.88<br>(-4.87; -0.88)<br>$p = .005$ | | |
| **8** | 6.61<br>(4.36; 8.85)<br>$p < .001$ | 5.28<br>(3.12; 7.44)<br>$p < .001$ | 1.27<br>(-1.23; 3.76)<br>$p = .319$ | 4.14<br>(2.26; 6.03)<br>$p < .001$ | |
| **12** | 12.57<br>(10.15; 15.00)<br>$p < .001$ | 11.24<br>(8.87; 13.62)<br>$p < .001$ | 7.23<br>(4.57; 9.89)<br>$p < .001$ | 10.11<br>(7.94; 12.27)<br>$p < .001$ | 5.96<br>(4.61; 7.32)<br>$p < .001$ |

**C01**

| Day | 0 | 2 | 3 | 4 | 8 |
|---|---|---|---|---|---|
| **2** | 1.47<br>(0.57; 2.38)<br>$p = .002$ | | | | |
| **3** | 5.47<br>(4.29; 6.65)<br>$p < .001$ | 4.00<br>(2.53; 5.48)<br>$p < .001$ | | | |
| **4** | 2.91<br>(1.49; 4.33)<br>$p < .001$ | 1.44<br>(0.22; 2.66)<br>$p = .021$ | -2.56<br>(-4.38; -0.74)<br>$p = .006$ | | |
| **8** | 6.46<br>(4.43; 8.50)<br>$p < .001$ | 4.99<br>(3.05; 6.94)<br>$p < .001$ | 0.99<br>(-1.32; 3.30)<br>$p = .399$ | 3.55<br>(1.93; 5.17)<br>$p < .001$ | |
| **12** | 10.06<br>(7.76; 12.35)<br>$p < .001$ | 8.59<br>(6.36; 10.82)<br>$p < .001$ | 4.59<br>(2.05; 7.12)<br>$p < .001$ | 7.14<br>(5.15; 9.14)<br>$p < .001$ | 3.59<br>(2.30; 4.89)<br>$p < .001$ |

**C05**

| Day | 0 | 2 | 3 | 4 | 8 |
|---|---|---|---|---|---|
| **2** | 2.85<br>(2.00; 3.70)<br>$p < .001$ | | | | |
| **3** | 7.35<br>(6.17; 8.53)<br>$p < .001$ | 4.50<br>(3.06; 5.94)<br>$p < .001$ | | | |
| **4** | 5.19<br>(3.77; 6.61)<br>$p < .001$ | 2.34<br>(1.16; 3.52)<br>$p < .001$ | -2.16<br>(-3.98; -0.34)<br>$p = .020$ | | |
| **8** | 10.14<br>(8.11; 12.17)<br>$p < .001$ | 7.29<br>(5.37; 9.20)<br>$p < .001$ | 2.79<br>(0.48; 5.10)<br>$p < .018$ | 4.95<br>(3.33; 6.57)<br>$p < .001$ | |
| **12** | 14.62<br>(12.39; 16.85)<br>$p < .001$ | 11.77<br>(9.62; 13.91)<br>$p < .001$ | 7.27<br>(4.79; 9.75)<br>$p < .001$ | 9.43<br>(7.50; 11.35)<br>$p < .001$ | 4.48<br>(3.30; 5.67)<br>$p < .001$ |

**C1**

| Day | 0 | 2 | 3 | 4 | 8 |
|---|---|---|---|---|---|
| **2** | 3.91<br>(3.07; 4.76)<br>$p < .001$ | | | | |
| **3** | 8.11<br>(6.93; 9.29)<br>$p < .001$ | 4.19<br>(2.75; 5.64)<br>$p < .001$ | | | |
| **4** | 6.69<br>(5.27; 8.11)<br>$p < .001$ | 2.78<br>(1.60; 3.96)<br>$p < .001$ | -1.41<br>(-3.24; 0.41)<br>$p = .128$ | | |
| **8** | 11.54<br>(9.51; 13.57)<br>$p < .001$ | 7.63<br>(5.71; 9.54)<br>$p < .001$ | 3.43<br>(1.12; 5.74)<br>$p = .004$ | 4.85<br>(3.23; 6.47)<br>$p < .001$ | |
| **12** | 18.04<br>(15.80; 20.27)<br>$p < .001$ | 14.12<br>(11.98; 16.27)<br>$p < .001$ | 9.93<br>(7.45; 12.41)<br>$p < .001$ | 11.34<br>(9.42; 13.27)<br>$p < .001$ | 6.50<br>(5.31; 7.68)<br>$p < .001$ |

**C10**

| Day | 0 | 2 | 3 | 4 | 8 |
|---|---|---|---|---|---|
| **2** | 6.56<br>(5.57; 7.54)<br>$p < .001$ | | | | |
| **3** | 7.61<br>(6.32; 8.90)<br>$p < .001$ | 1.05<br>(-0.56; 2.66)<br>$p = .200$ | | | |
| **4** | 13.97<br>(12.55; 15.39)<br>$p < .001$ | 7.41<br>(6.13; 8.70)<br>$p < .001$ | 6.36<br>(4.47; 8.26)<br>$p < .001$ | | |
| **8** | 21.32<br>(19.23; 23.40)<br>$p < .001$ | 14.76<br>(12.73; 16.79)<br>$p < .001$ | 13.71<br>(11.30; 16.12)<br>$p < .001$ | 7.34<br>(5.66; 9.03)<br>$p < .001$ | |
| **12** | 43.02<br>(40.78; 45.28)<br>$p < .001$ | 36.47<br>(34.26; 38.68)<br>$p < .001$ | 35.42<br>(32.88; 37.96)<br>$p < .001$ | 29.05<br>(27.12; 30.98)<br>$p < .001$ | 21.71<br>(20.44; 22.98)<br>$p < .001$ |

**Table S3. Descriptive statistics of the differences in estimated marginal means of accumulated GAG loss between different groups at several time points in the pilot study.** The accumulated GAG loss is shown in the Table S1. The comparisons are shown between different groups on days

0, 2, 3, 4, 8, and 12 with 95% confidence intervals and the associated $p$-value. The $p$-values smaller than 0.05 and 0.01 are shown with light yellow and light red, respectively. The difference is shown as the mean difference in the total GAG loss in the specified group in row minus the group in column.

| Group | Day 0 Group CTRL | Day 0 Group C01 | Day 0 Group C05 | Day 0 Group C1 | Day 2 Group CTRL | Day 2 Group C01 | Day 2 Group C05 | Day 2 Group C1 |
|---|---|---|---|---|---|---|---|---|
| C01 | 0.14<br>(-2.67; 2.95)<br>$p$ = .923 | | | | 0.28<br>(-2.72; 3.28)<br>$p$ = .853 | | | |
| C05 | 0.28<br>(-2.53; 3.09)<br>$p$ = .843 | 0.14<br>(-2.67; 2.95)<br>$p$ = .920 | | | 1.80<br>(-1.18; 4.79)<br>$p$ = .232 | 1.53<br>(-1.42; 4.47)<br>$p$ = .306 | | |
| C1 | 0.20<br>(-2.61; 3.01)<br>$p$ = .886 | 0.07<br>(-2.74; 2.88)<br>$p$ = .963 | -0.08<br>(-2.89; 2.73)<br>$p$ = .957 | | 2.79<br>(-0.19; 5.77)<br>$p$ = .066 | 2.51<br>(-0.44; 5.45)<br>$p$ = .094 | 0.98<br>(-1.95; 3.91)<br>$p$ = .506 | |
| C10 | 2.65<br>(-0.23; 5.52)<br>$p$ = .070 | 2.51<br>(-0.36; 5.39)<br>$p$ = .086 | 2.37<br>(-0.50; 5.24)<br>$p$ = .104 | 2.45<br>(-0.43; 5.32)<br>$p$ = .094 | 7.88<br>(4.81; 10.95)<br>$p$ < .001 | 7.60<br>(4.56; 10.64)<br>$p$ < .001 | 6.08<br>(3.05; 9.10)<br>$p$ < .001 | 5.09<br>(2.07; 8.12)<br>$p$ = .001 |

| Group | Day 3 Group CTRL | Day 3 Group C01 | Day 3 Group C05 | Day 3 Group C1 | Day 4 Group CTRL | Day 4 Group C01 | Day 4 Group C05 | Day 4 Group C1 |
|---|---|---|---|---|---|---|---|---|
| C01 | 0.27<br>(-2.77; 3.30)<br>$p$ = .862 | | | | 0.59<br>(-2.64; 3.81)<br>$p$ = .720 | | | |
| C05 | 2.29<br>(-0.75; 5.32)<br>$p$ = .138 | 2.02<br>(-1.01; 5.05)<br>$p$ = .189 | | | 3.01<br>(-0.22; 6.23)<br>$p$ = .067 | 2.42<br>(-0.70; 5.55)<br>$p$ = .127 | | |
| C1 | 2.97<br>(-0.07; 6.00)<br>$p$ = .055 | 2.70<br>(-0.33; 5.73)<br>$p$ = .080 | 0.68<br>(-2.35; 3.71)<br>$p$ = .657 | | 4.43<br>(1.20; 7.66)<br>$p$ = .008 | 3.85<br>(0.72; 6.97)<br>$p$ = .016 | 1.42<br>(-1.70; 4.55)<br>$p$ = .368 | |
| C10 | 4.92<br>(1.79; 8.04)<br>$p$ = .002 | 4.65<br>(1.52; 7.77)<br>$p$ = .004 | 2.63<br>(-0.50; 5.75)<br>$p$ = .098 | 1.95<br>(-1.18; 5.07)<br>$p$ = .219 | 14.16<br>(10.89; 17.43)<br>$p$ < .001 | 13.57<br>(10.40; 16.74)<br>$p$ < .001 | 11.15<br>(7.98; 14.32)<br>$p$ < .001 | 9.73<br>(6.56; 12.90)<br>$p$ < .001 |

| Group | Day 8 Group CTRL | Day 8 Group C01 | Day 8 Group C05 | Day 8 Group C1 | Day 12 Group CTRL | Day 12 Group C01 | Day 12 Group C05 | Day 12 Group C1 |
|---|---|---|---|---|---|---|---|---|
| C01 | -0.01<br>(-3.54; 3.53)<br>$p$ = .997 | | | | -2.38<br>(-6.05; 1.29)<br>$p$ = .202 | | | |
| C05 | 3.81<br>(0.27; 7.35)<br>$p$ = .035 | 3.82<br>(0.41; 7.23)<br>$p$ = .028 | | | 2.33<br>(-1.31; 5.96)<br>$p$ = .207 | 4.71<br>(1.16; 8.25)<br>$p$ = .010 | | |
| C1 | 5.14<br>(1.60; 8.67)<br>$p$ = .005 | 5.14<br>(1.74; 8.55)<br>$p$ = .003 | 1.33<br>(-2.08; 4.73)<br>$p$ = .442 | | 5.67<br>(2.04; 9.30)<br>$p$ = .003 | 8.05<br>(4.50; 11.59)<br>$p$ < .001 | 3.34<br>(-0.17; 6.85)<br>$p$ = .062 | |
| C10 | 17.36<br>(13.77; 20.95)<br>$p$ < .001 | 17.37<br>(13.90; 20.83)<br>$p$ < .001 | 13.55<br>(10.08; 17.01)<br>$p$ < .001 | 12.22<br>(8.76; 15.69)<br>$p$ < .001 | 33.10<br>(29.45; 36.76)<br>$p$ < .001 | 35.48<br>(31.91; 39.05)<br>$p$ < .001 | 30.78<br>(27.24; 34.31)<br>$p$ < .001 | 27.44<br>(23.91; 30.97)<br>$p$ < .001 |

#### S2.2 Aggrecan biosynthesis is lowered with high IL-1α concentration

In the pilot study, the biosynthesis rate of aggrecan normalized with the amount of DNA in cartilage plugs was significantly decreased with high concentrations of IL-1α (Fig. S3, Table S4). The analysis of time-dependent changes in  biosynthesis was not applicable in the pilot study since we used two different animals for different time points (Table S5). Aggrecan biosynthesis rate was suppressed with 10 ng/ml of IL-1α (median rate 8 pmol/h/μg) over a 12-day culture compared to CTRL (58 pmol/h/μg, $p$ = 0.020) and concentrations of 0.5 ng/ml (79 pmol/h/μg, $p$ = 0.001) and 1 ng/ml (86 pmol/h/μg, $p$ = 0.001; Fig. S3, Table S6). The notifiable increase in cell death under aggressive cytokine challenge[1,2] may especially affect the results obtained with 10 ng/ml and 12-day culture. This further highlights the suitability of 1 ng/ml for our study since 10 ng/ml might greatly hamper the aggrecan biosynthesis rate of cartilage. However, we found no significant differences in biosynthesis between different cytokine concentrations over a 7–day culture (Table S6). In Tables S4-S6, the IL-1α-challenge-only groups with concentrations of 0.1 ng/ml, 0.5 ng/ml, 1 ng/ml, and 10 ng/ml are marked as C01, C05, C1, and C10, respectively.

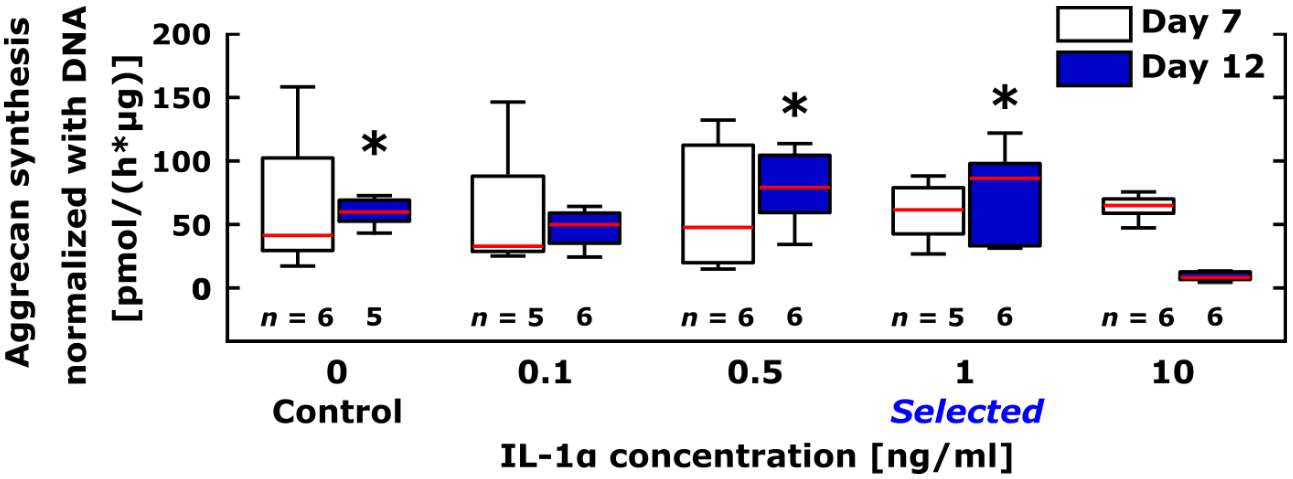


**Fig. S3. Aggrecan biosynthesis in the pilot study with different concentrations of interleukin (IL)-1α to find a suitable concentration for the rest of the study.** We observed a significant decrease in aggrecan biosynthesis rate after a 12-day culture with a concentration of 10 ng/ml of IL-1α compared to the control and cytokine concentrations of 0.5 ng/ml and 1 ng/ml (* = significant difference compared to the day 12 biosynthesis rate of the 10 ng/ml group, $p < 0.05$, linear mixed effects model). The IL-1α concentration of 1 ng/ml was chosen for the rest of the study since that concentration did not suppress aggrecan biosynthesis as aggressively as the concentration of 10 ng/ml. The data without location-matching of the samples was collected at different time points (days 7 and 12) from two animals (one animal for each time point, the amount of plugs *n* designated for each group is shown below the boxplots).

**Table S4. Aggrecan biosynthesis rates in the pilot study.** The results are shown as the estimated marginal mean values (pmol/h/μg) with standard error from the linear mixed effect models. *N* is the number of animals and *n* the total amount of plugs combined from those animals. The total amount of animals in the pilot study was 2.

| Group | Day 7, $N = 1$ | Day 12, $N = 1$ |
|---|---|---|
| CTRL | 65.01 (32.06) $n = 6$ | 57.68 (32.63) $n = 5$ |
| C01 | 58.37 (32.63) $n = 5$ | 47.07 (32.06) $n = 6$ |
| C05 | 62.44 (32.06) $n = 6$ | 78.32 (32.06) $n = 6$ |
| C1 | 57.92 (32.63) $n = 5$ | 76.18 (32.06) $n = 6$ |
| C10 | 63.61 (32.06) $n = 6$ | 8.92 (32.06) $n = 6$ |

**Table S5. Descriptive statistics of the differences in estimated marginal means of aggrecan biosynthesis rate between different time points within groups in the pilot study.** The aggrecan biosynthesis rates are shown in the Table S4. The comparisons are shown between days 7 and 12 with 95% confidence intervals and the associated *p*-value. The difference is shown as the mean difference in the aggrecan biosynthesis (pmol/h/μg) in the specified group in row minus the group in column.

| Day | CTRL Day 7 | C01 Day 7 | C05 Day 7 | C1 Day 7 | C10 Day 7 |
|---|---|---|---|---|---|
| 12 | -7.32 (-230.41; 215.76) $p > .999$ | -11.30 (-234.39; 211.79) $p > .999$ | 15.88 (-220.65; 252.42) $p > .999$ | 18.26 (-204.83; 241.35) $p > .999$ | -54.68 (-291.21; 181.85) $p > .999$ |

**Table S6. Descriptive statistics of the differences in estimated marginal means of aggrecan biosynthesis between different groups at different time points in the pilot study.** The aggrecan biosynthesis rates are shown in the Table S4. The comparisons are shown between different groups on days 7 and 12 with 95% confidence intervals and the associated *p*-value. The *p*-values smaller than 0.05 and 0.01 are shown with light yellow and light red, respectively. The difference is shown as the mean difference in the aggrecan biosynthesis (pmol/h/µg) in the specified group in row minus the group in column.

| | Day 7 Group | | | | Day 12 Group | | | |
|---|---|---|---|---|---|---|---|---|
| **Group** | **CTRL** | **C01** | **C05** | **C1** | **CTRL** | **C01** | **C05** | **C1** |
| **C01** | **-6.64<br>(-47.24; 33.97)<br>*p* = .744** | | | | **-10.61<br>(-51.22; 29.99)<br>*p* = .601** | | | |
| **C05** | **-2.57<br>(-41.28; 36.15)<br>*p* = .894** | **4.07<br>(-36.53; 44.67)<br>*p* = .841** | | | **20.64<br>(-19.96; 61.24)<br>*p* = .312** | **31.25<br>(-7.46; 69.97)<br>*p* = .111** | | |
| **C1** | **-7.09<br>(-47.69; 33.52)<br>*p* = .727** | **-0.45<br>(-42.86; 41.96)<br>*p* = .983** | **-4.52<br>(-45.12; 36.08)<br>*p* = .824** | | **18.50<br>(-22.10; 59.10)<br>*p* = .364** | **29.11<br>(-9.60; 67.83)<br>*p* = .137** | **-2.14<br>(-40.85; 36.57)<br>*p* = .912** | |
| **C10** | **-1.40<br>(-40.11; 37.31)<br>*p* = .942** | **5.24<br>(-35.37; 45.84)<br>*p* = .796** | **1.17<br>(-37.55; 39.88)<br>*p* = .952** | **5.69<br>(-34.92; 46.23)<br>*p* = .779** | **-48.76<br>(-89.36; -8.16)<br>*p* = .020** | **-38.15<br>(-76.86; 0.57)<br>*p* = .053** | **-69.40<br>(-108.11; -30.69)<br>*p* = .001** | **-67.26<br>(-105.97; -28.55)<br>*p* = .001** |

#### S2.3 Cyclic loading does not decelerate GAG loss significantly in IL-1 inflamed cartilage – injuries increase the GAG loss even more

The accumulated GAG loss increased significantly in all the treated groups (Tables S7 and S8). The accumulated GAG loss in the IL-CL groups was lower (for example, day 12 average 25.7%) than in the IL group (33.1%) but this difference was non-significant throughout the 12-day treatment (day 12 $p = 0.877$; Table S9). According to the mixed effects model, both of these treatments were more devastating than mere injuries (INJ group day 12 average 27.1%; INJ *vs.* IL $p = 0.003$, INJ *vs.* IL-CL $p = 0.033$). Applying an injurious loading in combination with inflammation increased the accumulated GAG loss (INJ-IL group day 12 average 40.0%, INJ-IL-CL group day 12 average 29.8%; IL *vs.* INJ-IL $p < 0.001$, IL-CL *vs.* INJ-IL-CL $p = 0.016$). However, this synergistic degradation via injuries and IL-1 was slowed down with cyclic loading in terms of (non-normalized) accumulated GAG loss (INJ-IL *vs.* INJ-IL-CL, $p = 0.037$).

**Table S7. Accumulated glycosaminoglycan (GAG) loss.** The results are shown as the estimated marginal mean values of accumulated GAG loss (%) with standard error from the linear mixed effect models. *N* is the number of animals and *n* the total amount of plugs combined from those animals. The total amount of animals was six; four for groups without and two for groups with cyclic loading.

| | Day | | | | | | | | | |
|---|---|---|---|---|---|---|---|---|---|---|
| Group | 0 | 2 | 3 | 4 | 5 | 6 | 7 | 8 | 10 | 12 |
| CTRL | 5.60<br>(1.44)<br>*N* = 6<br>*n* = 48 | 9.57<br>(1.43)<br>*N* = 6<br>*n* = 48 | 10.91<br>(1.46)<br>*N* = 4<br>*n* = 12 | 12.62<br>(1.44)<br>*N* = 5<br>*n* = 30 | 13.80<br>(1.46)<br>*N* = 5<br>*n* = 18 | 15.20<br>(1.47)<br>*N* = 4<br>*n* = 18 | 16.58<br>(1.50)<br>*N* = 4<br>*n* = 12 | 18.06<br>(1.49)<br>*N* = 4<br>*n* = 18 | 20.87<br>(1.49)<br>*N* = 4<br>*n* = 18 | 22.79<br>(1.51)<br>*N* = 4<br>*n* = 12 |
| INJ | 5.31<br>(1.49)<br>*N* = 4<br>*n* = 30 | 10.72<br>(1.49)<br>*N* = 4<br>*n* = 28 | 12.48<br>(1.56)<br>*N* = 2<br>*n* = 5 | 14.63<br>(1.50)<br>*N* = 3<br>*n* = 18 | 15.66<br>(1.52)<br>*N* = 3<br>*n* = 12 | 17.91<br>(1.53)<br>*N* = 2<br>*n* = 12 | 19.15<br>(1.61)<br>*N* = 2<br>*n* = 6 | 20.76<br>(1.56)<br>*N* = 2<br>*n* = 12 | 23.80<br>(1.57)<br>*N* = 2<br>*n* = 12 | 26.34<br>(1.61)<br>*N* = 2<br>*n* = 6 |
| IL | 5.32<br>(1.49)<br>*N* = 4<br>*n* = 29 | 10.58<br>(1.49)<br>*N* = 4<br>*n* = 28 | 12.52<br>(1.55)<br>*N* = 2<br>*n* = 6 | 14.53<br>(1.50)<br>*N* = 3<br>*n* = 18 | 15.42<br>(1.52)<br>*N* = 3<br>*n* = 12 | 17.92<br>(1.53)<br>*N* = 2<br>*n* = 12 | 18.47<br>(1.61)<br>*N* = 2<br>*n* = 6 | 23.07<br>(1.56)<br>*N* = 2<br>*n* = 12 | 26.62<br>(1.57)<br>*N* = 2<br>*n* = 12 | 29.70<br>(1.61)<br>*N* = 2<br>*n* = 6 |
| INJ-IL | 6.28<br>(1.49)<br>*N* = 4<br>*n* = 28 | 12.93<br>(1.49)<br>*N* = 4<br>*n* = 28 | 14.56<br>(1.55)<br>*N* = 2<br>*n* = 6 | 16.93<br>(1.50)<br>*N* = 3<br>*n* = 17 | 18.44<br>(1.52)<br>*N* = 3<br>*n* = 12 | 21.42<br>(1.54)<br>*N* = 2<br>*n* = 11 | 23.41<br>(1.61)<br>*N* = 2<br>*n* = 6 | 25.38<br>(1.57)<br>*N* = 2<br>*n* = 11 | 30.38<br>(1.58)<br>*N* = 2<br>*n* = 11 | 36.08<br>(1.64)<br>*N* = 2<br>*n* = 5 |
| IL-CL | 6.97<br>(1.59)<br>*N* = 2<br>*n* = 18 | 11.26<br>(1.59)<br>*N* = 2<br>*n* = 18 | 13.45<br>(1.64)<br>*N* = 2<br>*n* = 6 | 15.62<br>(1.61)<br>*N* = 2<br>*n* = 12 | 17.76<br>(1.66)<br>*N* = 2<br>*n* = 6 | 17.87<br>(1.69)<br>*N* = 2<br>*n* = 6 | 21.90<br>(1.72)<br>*N* = 2<br>*n* = 6 | 21.94<br>(1.74)<br>*N* = 2<br>*n* = 6 | 25.95<br>(1.76)<br>*N* = 2<br>*n* = 6 | 29.48<br>(1.77)<br>*N* = 2<br>*n* = 6 |
| INJ-IL-CL | 7.53<br>(1.60)<br>*N* = 2<br>*n* = 17 | 11.65<br>(1.59)<br>*N* = 2<br>*n* = 17 | 14.48<br>(1.64)<br>*N* = 2<br>*n* = 6 | 15.92<br>(1.62)<br>*N* = 2<br>*n* = 10 | 17.64<br>(1.66)<br>*N* = 2<br>*n* = 6 | 18.32<br>(1.70)<br>*N* = 2<br>*n* = 6 | 21.34<br>(1.72)<br>*N* = 2<br>*n* = 6 | 24.04<br>(1.74)<br>*N* = 2<br>*n* = 6 | 28.83<br>(1.76)<br>*N* = 2<br>*n* = 6 | 32.95<br>(1.73)<br>*N* = 2<br>*n* = 6 |

**Table S8. Descriptive statistics of the differences in estimated marginal means of accumulated GAG loss between different time points within groups.** The accumulated GAG loss is shown in the Table S7. The comparisons are shown between days 0, 2, 3, 4, 8, and 12 with 95% confidence intervals and the associated *p*-value. The *p*-values smaller than 0.05 and 0.01 are shown with light yellow and light red, respectively. The difference is shown as the mean difference in the accumulated GAG loss in the specified group in row minus the group in column.

| | CTRL<br>Day | | | | | INJ<br>Day | | | | |
|---|---|---|---|---|---|---|---|---|---|---|
| Day | 0 | 2 | 3 | 4 | 8 | 0 | 2 | 3 | 4 | 8 |
| 2 | 3.97<br>(3.63; 4.31)<br>$p < .001$ | | | | | 5.41<br>(4.97; 5.84)<br>$p < .001$ | | | | |
| 3 | 5.31<br>(4.57; 6.04)<br>$p < .001$ | 1.34<br>(0.68; 2.00)<br>$p < .001$ | | | | 7.17<br>(6.07; 8.27)<br>$p < .001$ | 1.76<br>(0.74; 2.78)<br>$p = .001$ | | | |
| 4 | 7.02<br>(6.39; 7.65)<br>$p < .001$ | 3.05<br>(2.50; 3.61)<br>$p < .001$ | 1.71<br>(0.86; 2.56)<br>$p < .001$ | | | 9.31<br>(8.51; 10.11)<br>$p < .001$ | 3.91<br>(3.19; 4.62)<br>$p < .001$ | 2.14<br>(0.91; 3.37)<br>$p = .001$ | | |
| 8 | 12.46<br>(11.36; 13.55)<br>$p < .001$ | 8.49<br>(7.42; 9.56)<br>$p < .001$ | 7.15<br>(5.91; 8.38)<br>$p < .001$ | 5.44<br>(4.46; 6.42)<br>$p < .001$ | | 15.45<br>(14.09; 16.80)<br>$p < .001$ | 10.04<br>(8.71; 11.36)<br>$p < .001$ | 8.28<br>(6.62; 9.93)<br>$p < .001$ | 6.13<br>(4.93; 7.34)<br>$p < .001$ | |
| 12 | 17.19<br>(15.92; 18.45)<br>$p < .001$ | 13.22<br>(11.98; 14.46)<br>$p < .001$ | 11.88<br>(10.49; 13.26)<br>$p < .001$ | 10.17<br>(8.98; 11.35)<br>$p < .001$ | 4.73<br>(3.89; 5.57)<br>$p < .001$ | 21.02<br>(19.39; 22.65)<br>$p < .001$ | 15.62<br>(14.01; 17.23)<br>$p < .001$ | 13.85<br>(11.97; 15.74)<br>$p < .001$ | 11.71<br>(10.18; 13.25)<br>$p < .001$ | 5.58<br>(4.45; 6.71)<br>$p < .001$ |

| | IL<br>Day | | | | | INJ-IL<br>Day | | | | |
|---|---|---|---|---|---|---|---|---|---|---|
| Day | 0 | 2 | 3 | 4 | 8 | 0 | 2 | 3 | 4 | 8 |
| 2 | 5.26<br>(4.82; 5.70)<br>$p < .001$ | | | | | 6.65<br>(6.21; 7.09)<br>$p < .001$ | | | | |
| 3 | 7.20<br>(6.18; 8.22)<br>$p < .001$ | 1.94<br>(1.01; 2.87)<br>$p < .001$ | | | | 8.28<br>(7.26; 9.30)<br>$p < .001$ | 1.63<br>(0.70; 2.56)<br>$p = .001$ | | | |
| 4 | 9.22<br>(8.40; 10.03)<br>$p < .001$ | 3.96<br>(3.24; 4.68)<br>$p < .001$ | 2.01<br>(0.85; 3.18)<br>$p = .001$ | | | 10.65<br>(9.82; 11.48)<br>$p < .001$ | 4.00<br>(3.27; 4.73)<br>$p < .001$ | 2.37<br>(1.20; 3.55)<br>$p < .001$ | | |
| 8 | 17.76<br>(16.39; 19.12)<br>$p < .001$ | 12.49<br>(11.17; 13.82)<br>$p < .001$ | 10.55<br>(8.95; 12.15)<br>$p < .001$ | 8.54<br>(7.34; 9.74)<br>$p < .001$ | | 19.10<br>(17.68; 20.52)<br>$p < .001$ | 12.45<br>(11.07; 13.83)<br>$p < .001$ | 10.82<br>(9.18; 12.46)<br>$p < .001$ | 8.45<br>(7.19; 9.70)<br>$p < .001$ | |
| 12 | 24.39<br>(22.75; 26.03)<br>$p < .001$ | 19.13<br>(17.51; 20.74)<br>$p < .001$ | 17.19<br>(15.35; 19.03)<br>$p < .001$ | 15.17<br>(13.64; 16.71)<br>$p < .001$ | 6.63<br>(5.51; 7.76)<br>$p < .001$ | 29.80<br>(28.07; 31.52)<br>$p < .001$ | 23.15<br>(21.45; 24.85)<br>$p < .001$ | 21.52<br>(19.60; 23.44)<br>$p < .001$ | 19.14<br>(17.52; 20.77)<br>$p < .001$ | 10.70<br>(9.48; 11.91)<br>$p < .001$ |

| | IL-CL<br>Day | | | | | INJ-IL-CL<br>Day | | | | |
|---|---|---|---|---|---|---|---|---|---|---|
| Day | 0 | 2 | 3 | 4 | 8 | 0 | 2 | 3 | 4 | 8 |
| 2 | 4.29<br>(3.75; 4.84)<br>$p < .001$ | | | | | 4.13<br>(3.56; 4.69)<br>$p < .001$ | | | | |
| 3 | 6.48<br>(5.41; 7.55)<br>$p < .001$ | 2.19<br>(1.25; 3.12)<br>$p < .001$ | | | | 6.95<br>(5.81; 8.09)<br>$p < .001$ | 2.82<br>(1.81; 3.83)<br>$p = .001$ | | | |
| 4 | 8.65<br>(7.62; 9.69)<br>$p < .001$ | 4.36<br>(3.45; 5.27)<br>$p < .001$ | 2.17<br>(0.89; 3.46)<br>$p = .001$ | | | 8.40<br>(7.30; 9.49)<br>$p < .001$ | 4.27<br>(3.30; 5.23)<br>$p < .001$ | 1.45<br>(0.07; 2.82)<br>$p = .039$ | | |
| 8 | 14.98<br>(13.12; 16.84)<br>$p < .001$ | 10.68<br>(8.87; 12.50)<br>$p < .001$ | 8.50<br>(6.49; 10.51)<br>$p < .001$ | 6.32<br>(4.65; 8.00)<br>$p < .001$ | | 16.51<br>(14.63; 18.39)<br>$p < .001$ | 12.38<br>(10.56; 14.21)<br>$p < .001$ | 9.56<br>(7.52; 11.61)<br>$p < .001$ | 8.12<br>(6.36; 9.88)<br>$p < .001$ | |
| 12 | 22.51<br>(20.47; 24.56)<br>$p < .001$ | 18.22<br>(16.21; 20.23)<br>$p < .001$ | 16.03<br>(13.85; 18.21)<br>$p < .001$ | 13.86<br>(11.93; 15.78)<br>$p < .001$ | 7.53<br>(6.24; 8.83)<br>$p < .001$ | 25.42<br>(23.36; 27.48)<br>$p < .001$ | 21.29<br>(19.27; 23.31)<br>$p < .001$ | 18.47<br>(16.26; 20.68)<br>$p < .001$ | 17.02<br>(15.04; 19.01)<br>$p < .001$ | 8.91<br>(7.61; 10.20)<br>$p < .001$ |

**Table S9. Descriptive statistics of the differences in estimated marginal means of accumulated GAG loss between different groups at several time points.** The accumulated GAG loss is shown in the Table S7. The comparisons are shown between different groups on days 0, 2, 3, 4, 8, and 12 with 95% confidence intervals and the associated $p$-value. The $p$-values smaller than 0.05 and 0.01

are shown with light yellow and light red, respectively. The difference is shown as the mean difference in the accumulated GAG loss in the specified group in row minus the group in column.

| | Day 0 Group | | | | | Day 2 Group | | | | |
|---|---|---|---|---|---|---|---|---|---|---|
| Group | CTRL | INJ | IL | INJ-IL | IL-CL | CTRL | INJ | IL | INJ-IL | IL-CL |
| INJ | -0.29<br>(-1.65; 1.07)<br>*p* = .677 | | | | | 1.15<br>(-0.21; 2.52)<br>*p* = .097 | | | | |
| IL | -0.28<br>(-1.65; 1.08)<br>*p* = .682 | 0.00<br>(-1.40; 1.40)<br>*p* = .996 | | | | 1.01<br>(-0.36; 2.38)<br>*p* = .146 | -0.14<br>(-1.55; 1.26)<br>*p* = .841 | | | |
| INJ-IL | 0.68<br>(-0.70; 2.06)<br>*p* = .333 | 0.97<br>(-0.45; 2.38)<br>*p* = .178 | 0.96<br>(-0.45; 2.38)<br>*p* = .182 | | | 3.36<br>(1.99; 4.74)<br>*p* < .001 | 2.21<br>(0.80; 3.62)<br>*p* = .002 | 2.35<br>(0.94; 3.77)<br>*p* = .001 | | |
| IL-CL | 1.37<br>(-0.36; 3.09)<br>*p* = .121 | 1.65<br>(-0.47; 3.78)<br>*p* = .127 | 1.65<br>(-0.48; 3.78)<br>*p* = .128 | 0.69<br>(-1.45; 2.82)<br>*p* = .528 | | 1.69<br>(-0.03; 3.42)<br>*p* = .055 | 0.54<br>(-1.59; 2.67)<br>*p* = .618 | 0.68<br>(-1.45; 2.81)<br>*p* = .528 | -1.67<br>(-3.81; 0.47)<br>*p* = .125 | |
| INJ-IL-CL | 1.93<br>(0.19; 3.66)<br>*p* = .030 | 2.21<br>(0.08; 4.35)<br>*p* = .042 | 2.21<br>(0.08; 4.35)<br>*p* = .043 | 1.23<br>(-0.90; 3.39)<br>*p* = .253 | 0.56<br>(-1.25; 2.37)<br>*p* = .541 | 2.09<br>(0.36; 3.82)<br>*p* = .018 | 0.93<br>(-1.20; 3.07)<br>*p* = .389 | 1.08<br>(-1.06; 3.21)<br>*p* = .321 | -1.27<br>(-3.42; 0.87)<br>*p* = .242 | 0.40<br>(-1.41; 2.20)<br>*p* = .667 |
| | **Day 3 Group** | | | | | **Day 4 Group** | | | | |
| Group | CTRL | INJ | IL | INJ-IL | IL-CL | CTRL | INJ | IL | INJ-IL | IL-CL |
| INJ | 1.57<br>(-0.17; 3.32)<br>*p* = .077 | | | | | 2.01<br>(0.57; 3.44)<br>*p* = .006 | | | | |
| IL | 1.61<br>(-0.09; 3.31)<br>*p* = .063 | -0.04<br>(-1.91; 1.83)<br>*p* = .970 | | | | 1.91<br>(0.47; 3.35)<br>*p* = .009 | -0.09<br>(-1.59; 1.40)<br>*p* = .903 | | | |
| INJ-IL | 3.65<br>(1.94; 5.35)<br>*p* < .001 | 2.07<br>(0.20; 3.95)<br>*p* = .030 | 2.04<br>(0.21; 3.87)<br>*p* = .029 | | | 4.31<br>(2.86; 5.77)<br>*p* < .001 | 2.31<br>(0.80; 3.82)<br>*p* = .003 | 2.40<br>(0.90; 3.91)<br>*p* = .002 | | |
| IL-CL | 2.54<br>(0.58; 4.50)<br>*p* = .011 | 0.96<br>(-1.48; 3.40)<br>*p* = .438 | 0.93<br>(-1.48; 3.33)<br>*p* = .449 | -1.11<br>(-3.52; 1.30)<br>*p* = .365 | | 3.00<br>(1.17; 4.83)<br>*p* = .001 | 0.99<br>(-1.22; 3.21)<br>*p* = .378 | 1.09<br>(-1.13; 3.30)<br>*p* = .335 | -1.31<br>(-3.54; 0.92)<br>*p* = .247 | |
| INJ-IL-CL | 3.57<br>(1.59; 5.54)<br>*p* < .001 | 1.99<br>(-0.46; 4.44)<br>*p* = .110 | 1.96<br>(-0.46; 4.37)<br>*p* = .112 | -0.08<br>(-2.50; 2.34)<br>*p* = .948 | 1.03<br>(-1.08; 3.14)<br>*p* = .337 | 3.30<br>(1.44; 5.16)<br>*p* = .001 | 1.30<br>(-0.95; 3.54)<br>*p* = .256 | 1.39<br>(-0.85; 3.63)<br>*p* < .224 | -1.01<br>(-3.26; 1.24)<br>*p* = .378 | 0.30<br>(-1.67; 2.27)<br>*p* = .763 |
| | **Day 8 Group** | | | | | **Day 12 Group** | | | | |
| Group | CTRL | INJ | IL | INJ-IL | IL-CL | CTRL | INJ | IL | INJ-IL | IL-CL |
| INJ | 2.70<br>(0.90; 4.51)<br>*p* = .003 | | | | | 3.55<br>(1.51; 5.59)<br>*p* = .001 | | | | |
| IL | 5.01<br>(3.21; 6.82)<br>*p* < .001 | 2.31<br>(0.41; 4.22)<br>*p* = .018 | | | | 6.92<br>(4.87; 8.96)<br>*p* < .001 | 3.37<br>(1.16; 5.58)<br>*p* = .003 | | | |
| INJ-IL | 7.32<br>(5.48; 9.17)<br>*p* < .001 | 4.62<br>(2.68; 6.57)<br>*p* < .001 | 2.31<br>(0.36; 4.25)<br>*p* = .020 | | | 13.29<br>(11.17; 15.40)<br>*p* < .001 | 9.74<br>(7.46; 12.02)<br>*p* < .001 | 6.37<br>(4.09; 8.65)<br>*p* < .001 | | |
| IL-CL | 3.89<br>(1.55; 6.23)<br>*p* = .001 | 1.19<br>(-1.51; 3.88)<br>*p* = .387 | -1.13<br>(-3.82; 1.56)<br>*p* = .410 | -3.44<br>(-6.15; -0.72)<br>*p* = .013 | | 6.69<br>(4.21; 9.17)<br>*p* < .001 | 3.14<br>(0.26; 6.03)<br>*p* = .033 | -0.23<br>(-3.11; 2.66)<br>*p* = .877 | -6.60<br>(-9.53; -3.66)<br>*p* < .001 | |
| INJ-IL-CL | 5.98<br>(3.63; 8.33)<br>*p* < .001 | 3.28<br>(0.58; 5.98)<br>*p* = .017 | 0.97<br>(-1.73; 3.67)<br>*p* = .482 | -1.34<br>(-4.07; 1.39)<br>*p* = .334 | 2.10<br>(-0.55; 4.74)<br>*p* = .121 | 10.16<br>(7.67; 12.65)<br>*p* < .001 | 6.61<br>(3.72; 9.50)<br>*p* < .001 | 3.24<br>(0.35; 6.13)<br>*p* = .028 | -3.13<br>(-6.07; -0.19)<br>*p* = .037 | 3.47<br>(0.66; 6.27)<br>*p* = .016 |

In terms of normalized accumulated GAG loss (Table S10), the INJ-IL, IL-CL, and INJ-IL-CL groups released GAG significantly between the beginning of the treatment and day 12 (Table S11; INJ-IL $p < 0.001$, IL-CL $p < 0.001$, INJ-IL-CL $p < 0.001$). The plugs of the IL-CL group exhibited more normalized GAG loss compared to the plugs of the IL group on days 5, 7, 10, and 12 (for example, IL day 12 median fold change 1.33, IL-CL median fold change 1.47, $p$ = 0.005; Fig. S4egij, Table S12). Furthermore, from day 8 onwards, the INJ-IL-CL group exhibited significantly more normalized accumulated GAG loss compared to the INJ-IL group (Fig. S4hij, Table S12).

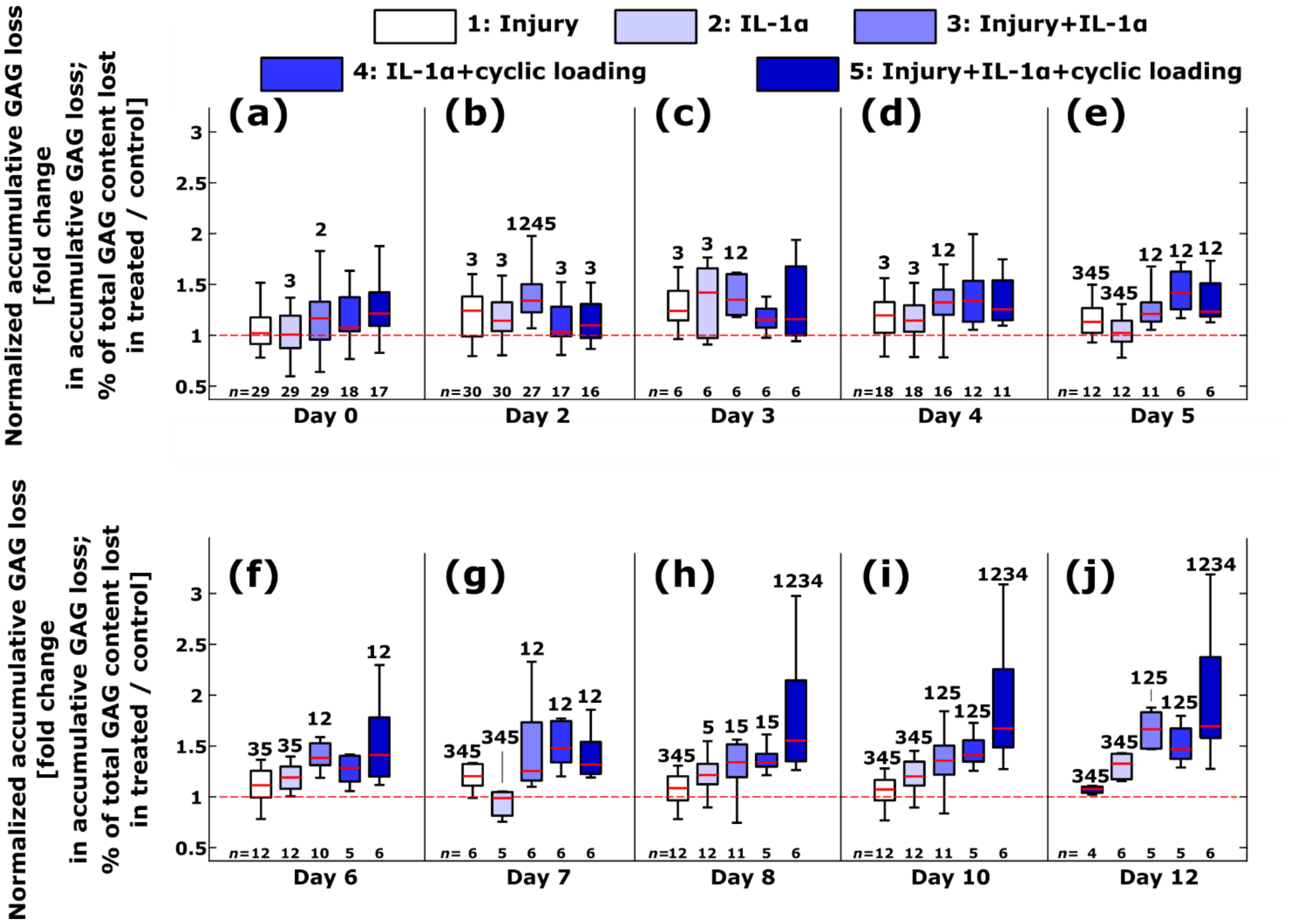


**Fig. S4. Normalized accumulated glycosaminoglycan (GAG) loss between different treatment groups at different time points**. Combination of injury and cytokines led to increased normalized accumulated GAG loss compared to either of these conditions alone. Moderate cyclic loading did not save cartilage from mechanical injury- and interleukin (IL)-1α-related disruption of proteoglycan matrix over the 12-day time period; such a loading scheme could, in turn, be disadvantageous. The amount of plugs $n$ designated for each group is shown on the bottom of each subpanel. The numbers above boxplots show statistically significant difference in normalized accumulated GAG loss compared to the indicated group(s) ($p < 0.05$; linear mixed effects model).

Normalized accumulated GAG loss depicts a fold change in accumulated GAG loss between a treated plug and that of an animal- and location-matched control.

**Table S10. Normalized accumulated glycosaminoglycan (GAG) loss.** The results are shown as the estimated marginal mean values (unitless; fold change in accumulated GAG loss in treated plugs versus in the control plugs) with standard error from the linear mixed effect models. *N* is the number of animals and *n* the total amount of plugs combined from those animals. The total amount of animals was six; four for groups without and two for groups with cyclic loading.

| | Day | | | | | | | | | |
|---|---|---|---|---|---|---|---|---|---|---|
| **Group** | **0** | **2** | **3** | **4** | **5** | **6** | **7** | **8** | **10** | **12** |
| **INJ** | 1.09<br>(0.06)<br>*N* = 4<br>*n* = 29 | 1.23<br>(0.06)<br>*N* = 4<br>*n* = 30 | 1.25<br>(0.07)<br>*N* = 2<br>*n* = 6 | 1.19<br>(0.06)<br>*N* = 3<br>*n* = 18 | 1.16<br>(0.07)<br>*N* = 3<br>*n* = 12 | 1.17<br>(0.07)<br>*N* = 2<br>*n* = 12 | 1.11<br>(0.08)<br>*N* = 2<br>*n* = 6 | 1.13<br>(0.07)<br>*N* = 2<br>*n* = 12 | 1.11<br>(0.08)<br>*N* = 2<br>*n* = 12 | 1.11<br>(0.09)<br>*N* = 2<br>*n* = 4 |
| **IL** | 1.06<br>(0.06)<br>*N* = 4<br>*n* = 29 | 1.20<br>(0.06)<br>*N* = 4<br>*n* = 30 | 1.22<br>(0.07)<br>*N* = 2<br>*n* = 6 | 1.19<br>(0.06)<br>*N* = 3<br>*n* = 18 | 1.17<br>(0.07)<br>*N* = 3<br>*n* = 12 | 1.16<br>(0.07)<br>*N* = 2<br>*n* = 12 | 1.12<br>(0.08)<br>*N* = 2<br>*n* = 5 | 1.20<br>(0.07)<br>*N* = 2<br>*n* = 12 | 1.18<br>(0.08)<br>*N* = 2<br>*n* = 12 | 1.19<br>(0.08)<br>*N* = 2<br>*n* = 6 |
| **INJ-IL** | 1.23<br>(0.06)<br>*N* = 4<br>*n* = 29 | 1.44<br>(0.06)<br>*N* = 4<br>*n* = 27 | 1.43<br>(0.07)<br>*N* = 2<br>*n* = 6 | 1.41<br>(0.06)<br>*N* = 3<br>*n* = 16 | 1.41<br>(0.07)<br>*N* = 3<br>*n* = 11 | 1.40<br>(0.07)<br>*N* = 2<br>*n* = 10 | 1.44<br>(0.08)<br>*N* = 2<br>*n* = 6 | 1.37<br>(0.08)<br>*N* = 2<br>*n* = 11 | 1.41<br>(0.08)<br>*N* = 2<br>*n* = 11 | 1.52<br>(0.09)<br>*N* = 2<br>*n* = 5 |
| **IL-CL** | 1.17<br>(0.08)<br>*N* = 2<br>*n* = 18 | 1.17<br>(0.08)<br>*N* = 2<br>*n* = 17 | 1.23<br>(0.09)<br>*N* = 2<br>*n* = 6 | 1.36<br>(0.08)<br>*N* = 2<br>*n* = 12 | 1.44<br>(0.09)<br>*N* = 2<br>*n* = 6 | 1.36<br>(0.10)<br>*N* = 2<br>*n* = 5 | 1.51<br>(0.10)<br>*N* = 2<br>*n* = 6 | 1.45<br>(0.11)<br>*N* = 2<br>*n* = 5 | 1.54<br>(0.11)<br>*N* = 2<br>*n* = 5 | 1.60<br>(0.11)<br>*N* = 2<br>*n* = 5 |
| **INJ-IL-CL** | 1.25<br>(0.08)<br>*N* = 2<br>*n* = 17 | 1.22<br>(0.08)<br>*N* = 2<br>*n* = 16 | 1.33<br>(0.09)<br>*N* = 2<br>*n* = 6 | 1.41<br>(0.08)<br>*N* = 2<br>*n* = 11 | 1.46<br>(0.09)<br>*N* = 2<br>*n* = 6 | 1.45<br>(0.09)<br>*N* = 2<br>*n* = 6 | 1.52<br>(0.10)<br>*N* = 2<br>*n* = 6 | 1.73<br>(0.10)<br>*N* = 2<br>*n* = 6 | 1.84<br>(0.10)<br>*N* = 2<br>*n* = 6 | 1.90<br>(0.11)<br>*N* = 2<br>*n* = 6 |

**Table S11. Descriptive statistics of the differences in estimated marginal means of normalized accumulated GAG loss between different time points within groups.** The normalized accumulated GAG loss is shown in the Table S10. The comparisons are shown between days 0, 2, 3, 4, 8, and 12 with 95% confidence intervals and the associated *p*-value. The *p*-values smaller than 0.05 and 0.01 are shown with light yellow and light red, respectively. The difference is shown as the mean difference in the normalized accumulated GAG loss in the specified group in row minus the group in column.

**INJ**

| Day | 0 | 2 | 3 | 4 | 8 |
|---|---|---|---|---|---|
| 2 | 0.13<br>(0.10; 0.17)<br>*p* < .001 | | | | |
| 3 | 0.15<br>(0.07; 0.24)<br>*p* < .001 | 0.02<br>(-0.06; 0.10)<br>*p* = .594 | | | |
| 4 | 0.10<br>(0.03; 0.17)<br>*p* = .004 | -0.03<br>(-0.09; 0.02)<br>*p* = .255 | -0.06<br>(-0.15; 0.04)<br>*p* = .263 | | |
| 8 | 0.03<br>(-0.08; 0.15)<br>*p* = .568 | -0.10<br>(-0.21; 0.02)<br>*p* = .088 | -0.12<br>(-0.26; 0.02)<br>*p* = .085 | -0.07<br>(-0.17; 0.04)<br>*p* = .213 | |
| 12 | 0.02<br>(-0.14; 0.17)<br>*p* = .818 | -0.12<br>(-0.27; 0.04)<br>*p* = .132 | -0.14<br>(-0.30; 0.03)<br>*p* = .111 | -0.08<br>(-0.23; 0.06)<br>*p* = .264 | -0.02<br>(-0.13; 0.09)<br>*p* = .769 |

**IL**

| Day | 0 | 2 | 3 | 4 | 8 |
|---|---|---|---|---|---|
| 2 | 0.14<br>(0.10; 0.17)<br>*p* < .001 | | | | |
| 3 | 0.16<br>(0.07; 0.24)<br>*p* < .001 | 0.02<br>(-0.06; 0.10)<br>*p* = .633 | | | |
| 4 | 0.13<br>(0.06; 0.20)<br>*p* < .001 | -0.01<br>(-0.07; 0.05)<br>*p* = .777 | -0.03<br>(-0.12; 0.07)<br>*p* = .578 | | |
| 8 | 0.14<br>(0.02; 0.25)<br>*p* = .024 | -0.00<br>(-0.12; 0.11)<br>*p* = .968 | -0.02<br>(-0.16; 0.12)<br>*p* = .761 | 0.01<br>(-0.10; 0.11)<br>*p* = .908 | |
| 12 | 0.13<br>(-0.01; 0.27)<br>*p* = .074 | -0.01<br>(-0.15; 0.13)<br>*p* = .907 | -0.03<br>(-0.19; 0.13)<br>*p* = .737 | 0.00<br>(-0.13; 0.13)<br>*p* = .999 | -0.01<br>(-0.10; 0.09)<br>*p* = .901 |

**INJ-IL**

| Day | 0 | 2 | 3 | 4 | 8 |
|---|---|---|---|---|---|
| 2 | 0.21<br>(0.18; 0.25)<br>*p* < .001 | | | | |
| 3 | 0.20<br>(0.12; 0.29)<br>*p* < .001 | -0.01<br>(-0.09; 0.06)<br>*p* = .742 | | | |
| 4 | 0.18<br>(0.11; 0.25)<br>*p* < .001 | -0.04<br>(-0.10; 0.03)<br>*p* = .260 | -0.02<br>(-0.12; 0.08)<br>*p* = .663 | | |
| 8 | 0.14<br>(0.02; 0.26)<br>*p* = .025 | -0.08<br>(-0.19; 0.04)<br>*p* = .215 | -0.06<br>(-0.20; 0.08)<br>*p* = .387 | -0.04<br>(-0.15; 0.07)<br>*p* = .460 | |
| 12 | 0.29<br>(0.14; 0.44)<br>*p* < .001 | 0.08<br>(-0.07; 0.22)<br>*p* = .316 | 0.09<br>(-0.08; 0.25)<br>*p* = .295 | 0.11<br>(-0.03; 0.25)<br>*p* = .124 | 0.15<br>(0.05; 0.25)<br>*p* = .004 |

**IL-CL**

| Day | 0 | 2 | 3 | 4 | 8 |
|---|---|---|---|---|---|
| 2 | -0.00<br>(-0.05; 0.04)<br>*p* = .891 | | | | |
| 3 | 0.06<br>(-0.03; 0.15)<br>*p* = .197 | 0.06<br>(-0.02; 0.14)<br>*p* = .117 | | | |
| 4 | 0.19<br>(0.10; 0.28)<br>*p* < .001 | 0.19<br>(0.12; 0.27)<br>*p* < .001 | 0.13<br>(0.02; 0.24)<br>*p* = .018 | | |
| 8 | 0.28<br>(0.11; 0.46)<br>*p* = .002 | 0.28<br>(0.11; 0.45)<br>*p* = .001 | 0.22<br>(0.04; 0.41)<br>*p* = .019 | 0.09<br>(-0.07; 0.25)<br>*p* = .255 | |
| 12 | 0.42<br>(0.23; 0.62)<br>*p* < .001 | 0.43<br>(0.23; 0.62)<br>*p* < .001 | 0.36<br>(0.16; 0.57)<br>*p* = .001 | 0.23<br>(0.05; 0.42)<br>*p* = .013 | 0.14<br>(0.02; 0.26)<br>*p* = .019 |

**INJ-IL-CL**

| Day | 0 | 2 | 3 | 4 | 8 |
|---|---|---|---|---|---|
| 2 | -0.04<br>(-0.09; 0.01)<br>*p* = .134 | | | | |
| 3 | 0.08<br>(-0.01; 0.17)<br>*p* = .091 | 0.12<br>(0.04; 0.20)<br>*p* = .005 | | | |
| 4 | 0.16<br>(0.07; 0.25)<br>*p* = .001 | 0.19<br>(0.11; 0.27)<br>*p* < .001 | 0.08<br>(-0.04; 0.19)<br>*p* = .178 | | |
| 8 | 0.48<br>(0.31; 0.64)<br>*p* < .001 | 0.51<br>(0.35; 0.67)<br>*p* < .001 | 0.40<br>(0.22; 0.57)<br>*p* < .001 | 0.32<br>(0.17; 0.47)<br>*p* < .001 | |
| 12 | 0.65<br>(0.47; 0.83)<br>*p* < .001 | 0.68<br>(0.51; 0.86)<br>*p* < .001 | 0.57<br>(0.37; 0.76)<br>*p* < .001 | 0.49<br>(0.32; 0.66)<br>*p* < .001 | 0.17<br>(0.06; 0.28)<br>*p* = .002 |

**Table S12. Descriptive statistics of the differences in estimated marginal means of normalized accumulated GAG loss between different groups at several time points.** The normalized accumulated GAG loss is shown in the Table S10. The comparisons are shown between different groups on days 0, 2, 3, 4, 8, and 12 with 95% confidence intervals and the associated *p*-value. The *p*-values smaller than 0.05 and 0.01 are shown with light yellow and light red, respectively. The difference is shown as the mean difference in the normalized accumulated GAG loss in the specified group in row minus the group in column.

| | Day 0 Group | | | | Day 2 Group | | | |
|---|---|---|---|---|---|---|---|---|
| **Group** | **INJ** | **IL** | **INJ-IL** | **IL-CL** | **INJ** | **IL** | **INJ-IL** | **IL-CL** |
| **IL** | -0.03 (-0.17; 0.11) $p = .654$ | | | | -0.03 (-0.17; 0.11) $p = .703$ | | | |
| **INJ-IL** | 0.13 (-0.01; 0.28) $p = .064$ | 0.17 (0.02; 0.31) $p = .022$ | | | 0.22 (0.07; 0.36) $p = .003$ | 0.24 (0.10; 0.38) $p = .001$ | | |
| **IL-CL** | 0.08 (-0.14; 0.30) $p = .426$ | 0.11 (-0.10; 0.33) $p = .273$ | -0.05 (-0.27; 0.17) $p = .612$ | | -0.05 (-0.27; 0.16) $p = .594$ | -0.03 (-0.24; 0.19) $p = .790$ | -0.27 (-0.49; -0.05) $p = .019$ | |
| **INJ-IL-CL** | 0.16 (-0.06; 0.38) $p = .134$ | 0.19 (-0.03; 0.41) $p = .078$ | 0.03 (-0.19; 0.24) $p = .795$ | 0.08 (-0.11; 0.26) $p = .399$ | -0.01 (-0.23; 0.21) $p = .921$ | 0.02 (-0.20; 0.23) $p = .866$ | -0.23 (-0.44; 0.01) $p = .044$ | 0.04 (-0.14; 0.23) $p = .633$ |

| | Day 3 Group | | | | Day 4 Group | | | |
|---|---|---|---|---|---|---|---|---|
| **Group** | **INJ** | **IL** | **INJ-IL** | **IL-CL** | **INJ** | **IL** | **INJ-IL** | **IL-CL** |
| **IL** | -0.03 (-0.20; 0.14) $p = .735$ | | | | -0.00 (-0.15; 0.15) $p = .980$ | | | |
| **INJ-IL** | 0.18 (0.01; 0.35) $p = .039$ | 0.21 (0.04; 0.38) $p = .017$ | | | 0.21 (0.07; 0.36) $p = .005$ | 0.22 (0.07; 0.36) $p = .005$ | | |
| **IL-CL** | -0.01 (-0.25; 0.22) $p = .905$ | 0.02 (-0.22; 0.25) $p = .885$ | -0.19 (-0.43; 0.04) $p = .096$ | | 0.17 (-0.05; 0.39) $p = .118$ | 0.17 (-0.05; 0.40) $p = .114$ | -0.04 (-0.26; 0.18) $p < .692$ | |
| **INJ-IL-CL** | 0.09 (-0.15; 0.32) $p = .454$ | 0.11 (-0.12; 0.35) $p = .315$ | -0.10 (-0.33; 0.14) $p = .397$ | 0.10 (-0.11; 0.30) $p = .344$ | 0.22 (-0.01; 0.44) $p = .056$ | 0.22 (-0.00; 0.44) $p = .054$ | 0.00 (-0.22; 0.23) $p = .984$ | 0.04 (-0.15; 0.24) $p = .654$ |

| | Day 8 Group | | | | Day 12 Group | | | |
|---|---|---|---|---|---|---|---|---|
| **Group** | **INJ** | **IL** | **INJ-IL** | **IL-CL** | **INJ** | **IL** | **INJ-IL** | **IL-CL** |
| **IL** | 0.07 (-0.11; 0.25) $p = .452$ | | | | 0.08 (-0.13; 0.29) $p = .463$ | | | |
| **INJ-IL** | 0.24 (0.05; 0.42) $p = .011$ | 0.17 (-0.02; 0.35) $p = .072$ | | | 0.41 (0.19; 0.62) $p < .001$ | 0.33 (0.11; 0.54) $p = .003$ | | |
| **IL-CL** | 0.33 (0.07; 0.59) $p = .015$ | 0.26 (-0.00; 0.52) $p = .051$ | 0.09 (-0.17; 0.35) $p = .491$ | | 0.49 (0.20; 0.77) $p = .001$ | 0.41 (0.13; 0.69) $p = .005$ | 0.08 (-0.20; 0.36) $p = .565$ | |
| **INJ-IL-CL** | 0.60 (0.35; 0.86) $p < .001$ | 0.53 (0.28; 0.79) $p < .001$ | 0.36 (0.11; 0.62) $p = .007$ | 0.27 (0.02; 0.53) $p = .037$ | 0.79 (0.51; 1.06) $p < .001$ | 0.71 (0.44; 0.98) $p < .001$ | 0.38 (0.11; 0.66) $p = .007$ | 0.30 (0.03; 0.58) $p = .031$ |

#### S2.4 Intact regions experienced less severe GAG loss in the superficial zone of inflamed cartilage after 12-day cyclic loading compared to samples without cyclic loading

For a more detailed figure of the optical density profiles on days 3 and 7, see Fig. S5.

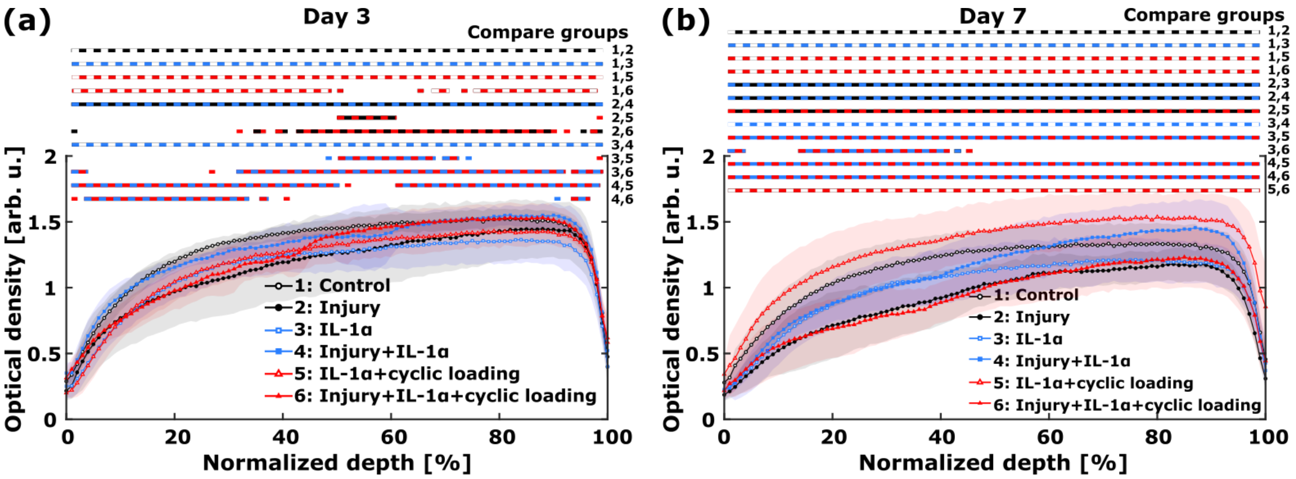


**Fig. S5. Optical density profiles on days 3 and 7 reveal depth-wise changes in localized glycosaminoglycan (GAG) content in response to different treatments.** The numbers show statistically significant differences ($p < 0.05$; linear mixed effects model; data shown as mean ± standard deviation) between groups at different normalized depths starting from the cartilage surface. **(a)** On day 3, the cyclically loaded groups showed marked loss of optical density especially in the superficial and transitional zones. **(b)** On day 7, the INJ-IL-CL group had significantly lower localized optical density throughout the whole cartilage depth compared to the INJ-IL group. The depth-wise GAG content was higher in the IL-CL group compared to the CTRL group. Abbreviations: CTRL = control, INJ-IL = injured and IL-1α-challenged, IL-CL = IL-1α-challenged and cyclically loaded, INJ-IL-CL = injured, IL-1α-challenged, and cyclically loaded.

Initially on the day 0, the depth-wise GAG contents away from lesions were higher in injured plugs compared to the uninjured plugs (Fig. S6a). Surprisingly, on day 5 (Fig. S6b), the INJ group exhibited the highest depth-wise GAG concentrations. On day 12 (Fig. S6d), all the treatment groups had similar depth-wise GAG content in intact regions as the CTRL group. Moreover, even on day 12, the cyclic loading still showed its beneficial features at least locally; the INJ-IL group showed significantly lower optical density than the INJ-IL-CL group in the transitional and deep zones ($p < 0.05$).

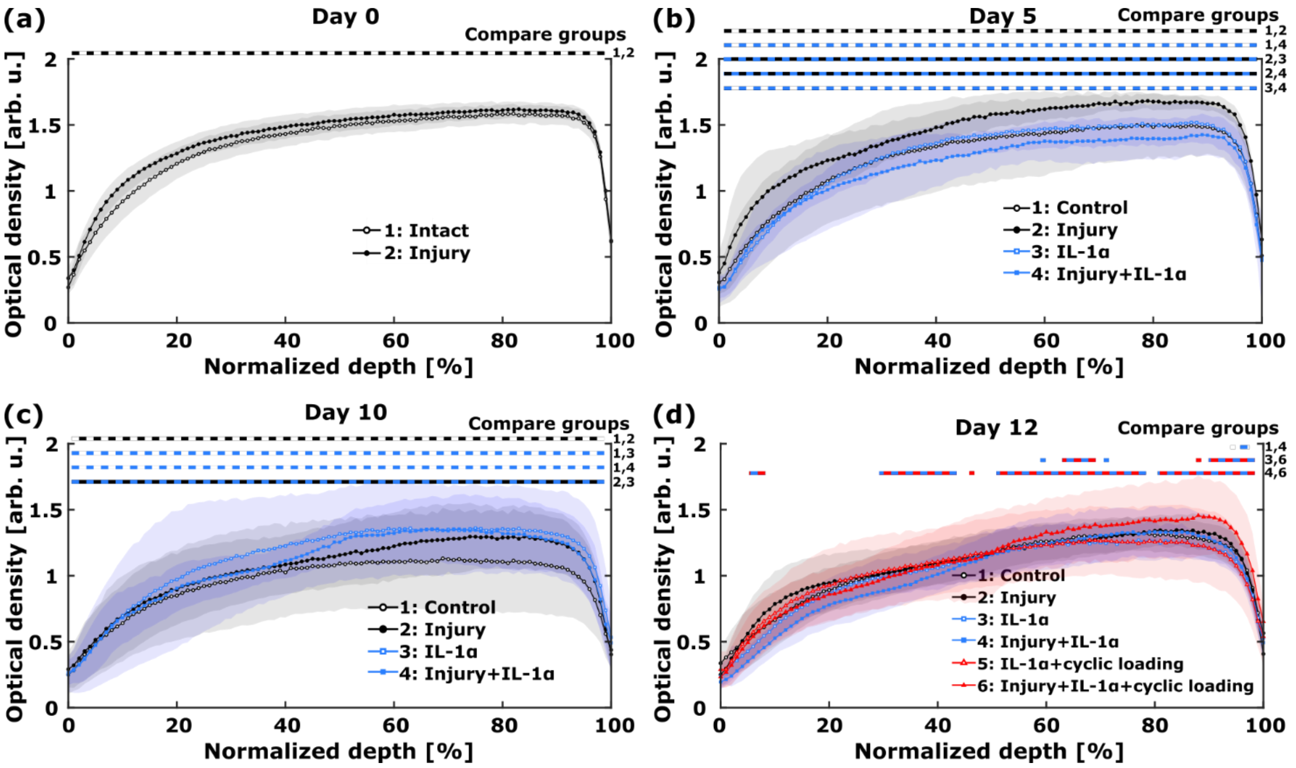


**Fig. S6. Optical density profiles on days 0, 5, 10, and 12 reveal depth-wise changes in localized glycosaminoglycan (GAG) content in response to different treatments. (a)** Initially, the injured samples exhibited higher optical densities away from lesions than the intact samples ($p < 0.05$; linear mixed effects model; the numbers show statistically significant differences between groups at different normalized depths starting from the cartilage surface; data shown as mean ± standard deviation). **(b)** On day 5 after initiation of the treatment, interestingly the INJ group exhibited the greatest levels of GAG away from the lesions. In contrast, the IL and INJ-IL groups showed marked depth-dependent depletion of GAG. **(c)** On day 10, some regions in the samples of the IL and INJ-IL groups could still exhibit higher levels of GAG than CTRL samples on average. **(d)** In general, on day 12, the optical density profiles indicate similar depth-dependent GAG concentrations in all the groups. However, we observed higher optical densities in the INJ-IL-CL group compared to the INJ-IL group in the deep regions of cartilage, suggesting a protective effect of cyclic loading in the deep zone still after a 12-day loading period. Abbreviations: CTRL = control, INJ = injury-only, IL = IL-1α-challenge-only, INJ-IL = injured and IL-1α-challenged, INJ-IL-CL = injured, IL-1α-challenged, and cyclically loaded.

In the main text Fig. 5 and here in the Figs. S5 and S6, the statistical significance between different groups at different cartilage depths was found by evaluating whether the mixed model-predicted mean optical density of one group falls into the 95% confidence interval of the mean optical density of another group. Here, we investigate the statistical significance with the help of confidence interval of difference. The groups to be compared are significantly different if the confidence interval of the difference of mean profiles does not include zero. As an example, on day 3, the INJ-IL-CL group had significantly higher optical density in the deep zone when compared to both INJ and IL groups (Fig. S7).

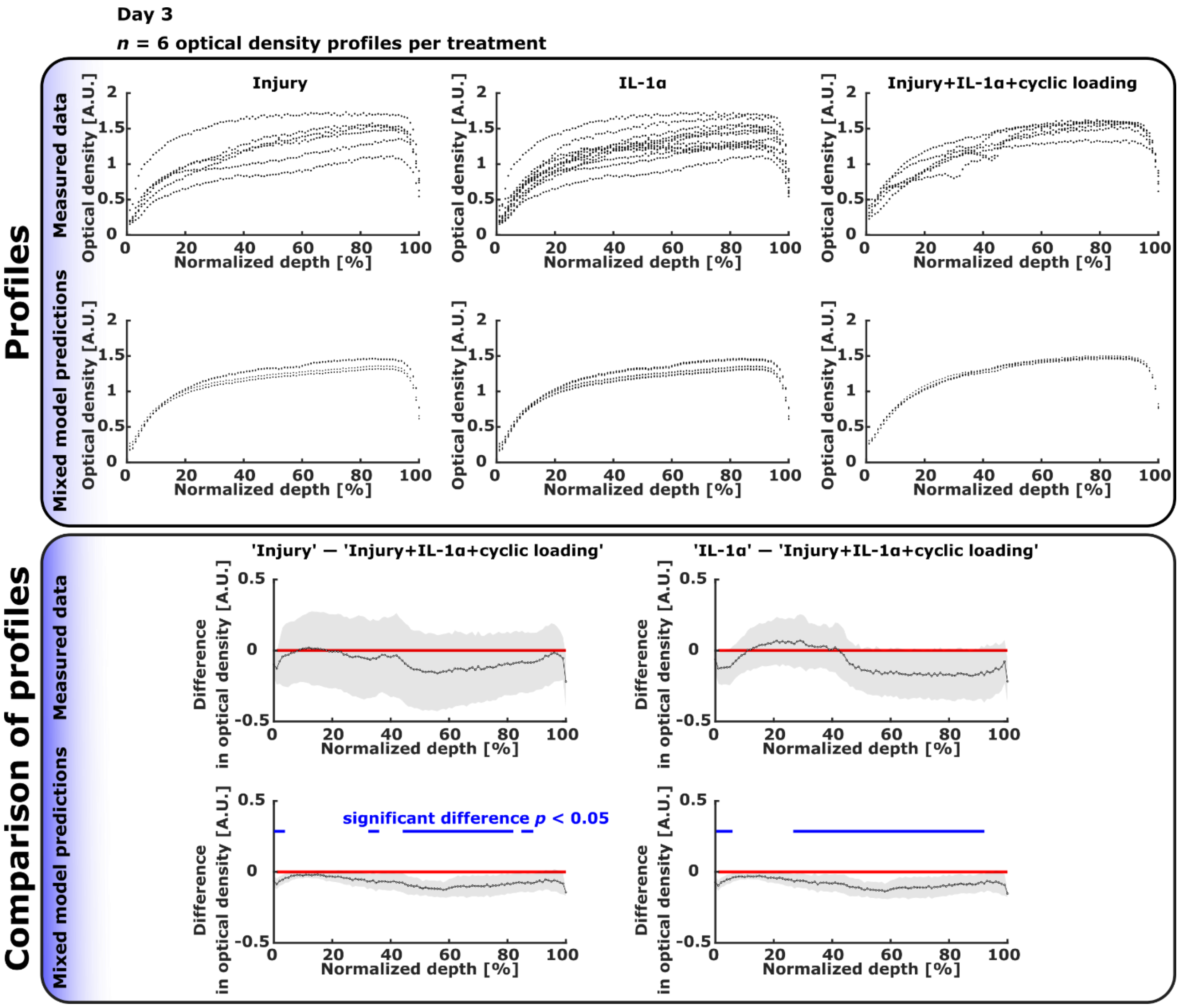

**Fig. S7. Further comparison of optical density profiles on day 3.** The top panel shows the measured optical density profiles and the corresponding predictions from the linear mixed effects model on day 3 after initiation of the treatment. The bottom panel shows the difference in mean optical densities of two specified groups (one group minus the other) with 95% confidence intervals calculated both from measured and predicted profiles. The compared groups were significantly different at depths where the confidence interval of the difference of mean profiles did not include zero (marked in blue, $p < 0.05$). The plugs in the INJ-IL-CL group had significantly higher optical density in the transitional/deep layers of cartilage compared to both INJ and IL groups. Abbreviations: INJ = injury-only, IL = IL-1α-challenge-only, INJ-IL-CL = injured, IL-1α-challenged, and cyclically loaded.

The GAG content near injuries was significantly lower compared to areas away from lesions in all injured groups at all time points (Fig. S8ac, Fig. S9aceg). After 12 days of culture, the cyclic loading still presented its cartilage health-promoting effects, at least locally; the average GAG concentration in intact regions was higher in cyclically loaded samples than in the INJ-IL group (Fig. S9h).

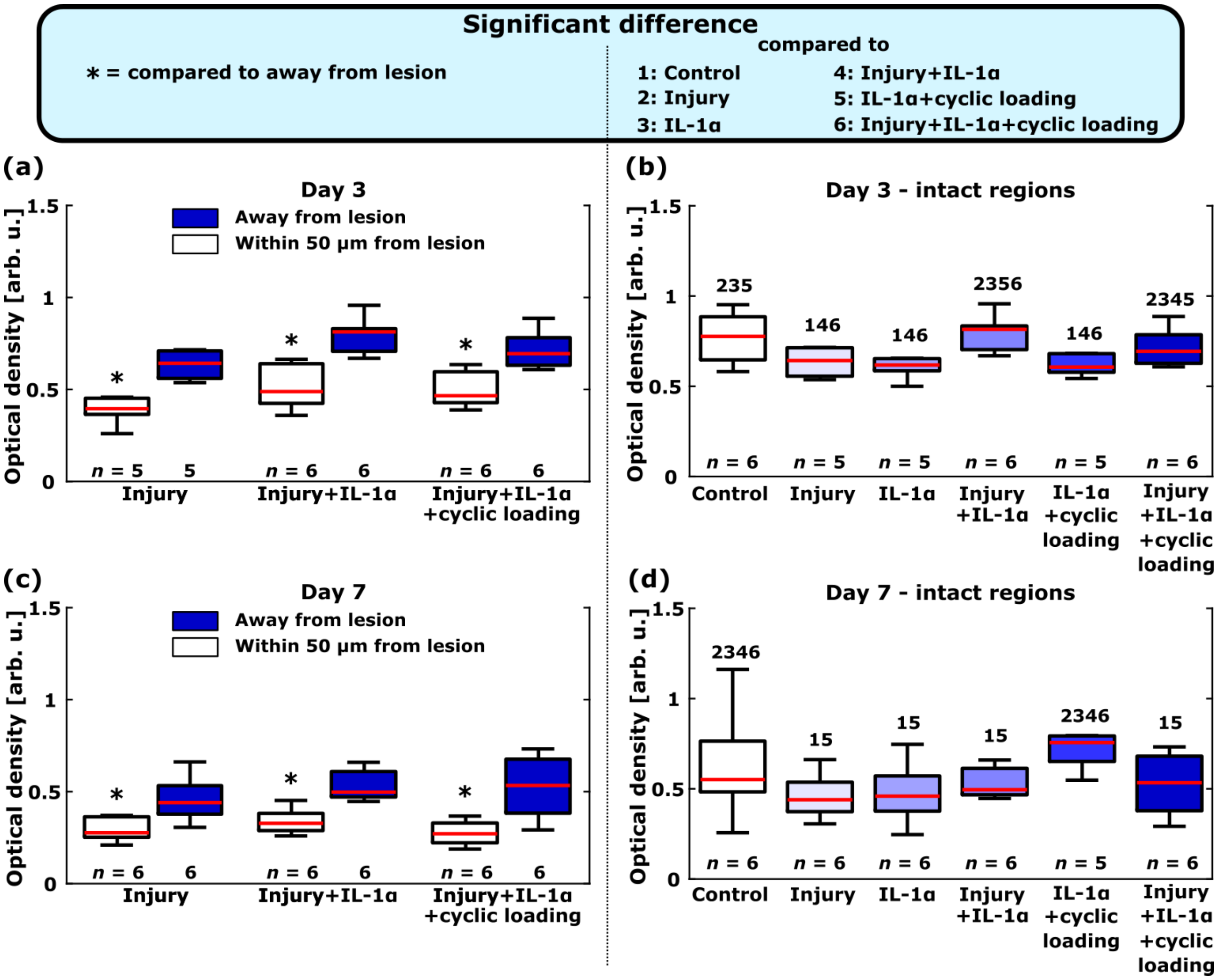


**Fig. S8. Localized glycosaminoglycan (GAG) content near and away from lesions on days 3 and 7.** On days 3 **(a)** and 7 **(c)** after beginning of the treatment, the localized average GAG content was significantly lower ($p < 0.05$; linear mixed effects model) near the lesions compared to away from lesions in all the injured groups. **(b)** The INJ-IL and INJ-IL-CL groups retained their GAG content better in regions away from lesions compared to the INJ, IL, and IL-CL groups. On day 7, **(d)** all the other groups except the IL-CL group lost their GAG content away from lesions below that of free-swelling control levels. The amount of plugs *n* designated for each group is shown on the bottom of subpanels. The numbers above boxplots (subpanels bd) show statistically significant difference in regional average optical density compared to the indicated group(s) ($p < 0.05$; linear mixed effects model). Abbreviations: INJ = injury-only, IL = IL-1α-challenge-only, INJ-IL =

injured and IL-1α-challenged, IL-CL = IL-1α-challenged and cyclically loaded, INJ-IL-CL = injured, IL-1α-challenged, and cyclically loaded.

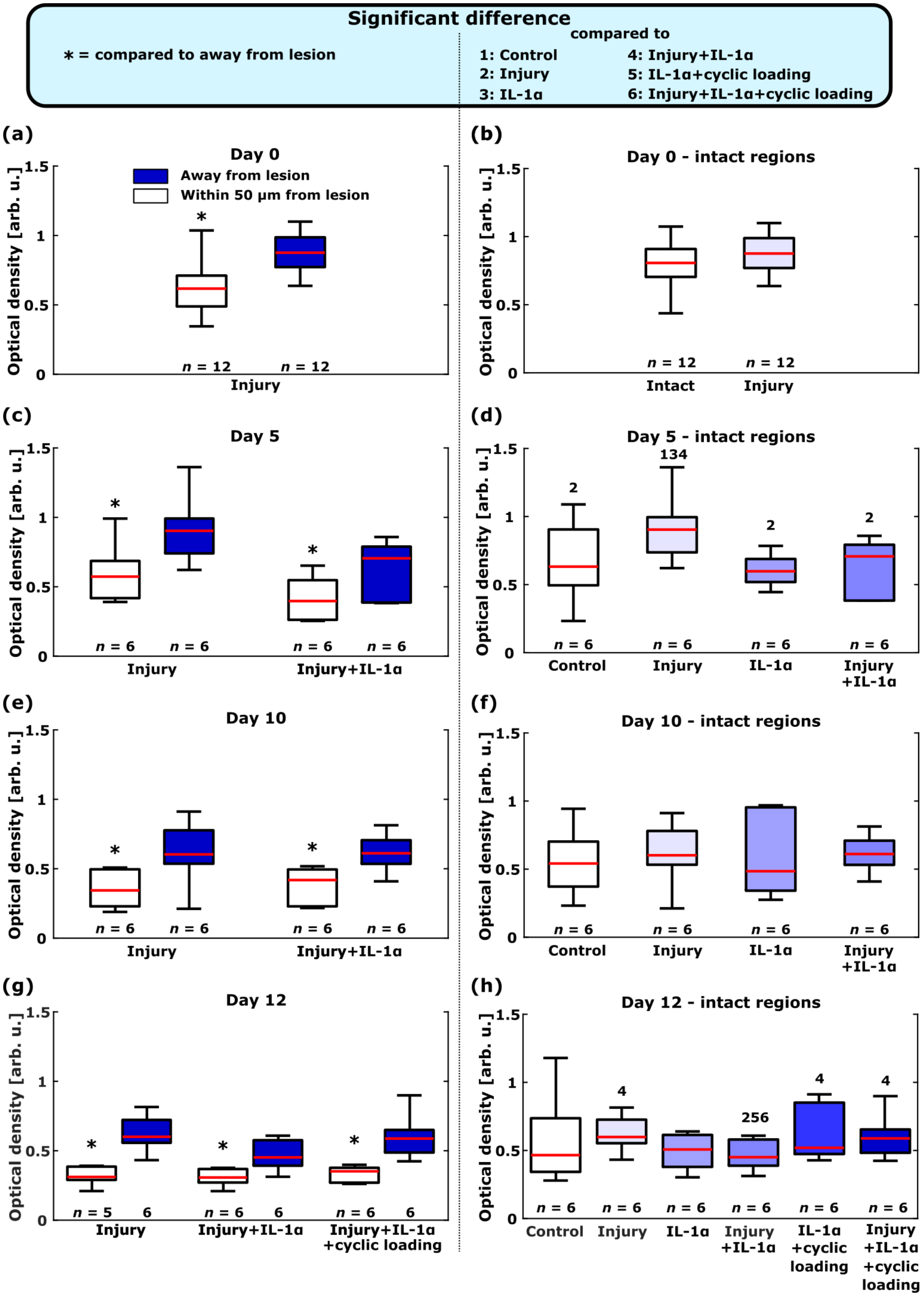

Significant difference
∗ = compared to away from lesion
compared to
1: Control
2: Injury
3: IL-1α
4: Injury+IL-1α
5: IL-1α+cyclic loading
6: Injury+IL-1α+cyclic loading
(a)
Day 0
Away from lesion
Within 50 μm from lesion
Optical density [arb. u.]
n = 12
n = 12
Injury
(b)
Day 0 - intact regions
Intact
Injury
(c)
Day 5
Injury
Injury+IL-1α
(d)
Day 5 - intact regions
134
Control
Injury
IL-1α
Injury +IL-1α
(e)
Day 10
(f)
Day 10 - intact regions
(g)
Day 12
Injury+IL-1α +cyclic loading
(h)
Day 12 - intact regions
256
IL-1α +cyclic loading
Injury +IL-1α +cyclic loading

**Fig. S9. Localized glycosaminoglycan (GAG) content near and away from lesions on days 0, 5, 10, and 12.** On the treatment initiation day, **(a)** the localized average GAG content was significantly lower ($p < 0.05$; linear mixed effects model) near the lesions compared to away from the lesions, **(b)** whereas the GAG concentrations were similar between intact plugs and those regions of injured plugs that were away from lesions. On day 5 after initiation of the treatment, **(c)** the GAG were retained worse near the lesions than away from them, and **(d)** the GAG concentration was surprisingly highest in the INJ group. On day 10, **(e)** the GAG loss was higher near the lesions compared to away from them, and **(f)** optical densities were similar away from lesions between all the groups. On day 12, **(g)** GAG was retained better away from the lesions than near them, and **(h)** cyclically loaded groups experienced less GAG loss in the superficial zone away from lesions compared to the INJ-IL group. The amount of plugs $n$ designated for each group is shown below the boxplots. The numbers above boxplots (subpanels bdfh) show statistically significant difference in regional average optical density compared to the indicated group(s) ($p < 0.05$; linear mixed effects model). Abbreviations: INJ = injury-only, INJ-IL = injured and IL-1α-challenged.

#### S2.5 Injury without exogenous inflammatory stress increases aggrecan biosynthesis transiently

Aggrecan biosynthesis rate (Table S13) decreased significantly over time in the IL group (Table S14). However, biosynthesis increased in the INJ group on days 7 and 10 compared to the INJ-IL group (Fig. S10cd, Table S15). Moreover, cyclic loading increased aggrecan biosynthesis but only after 12 days of loading (Fig. S10e).

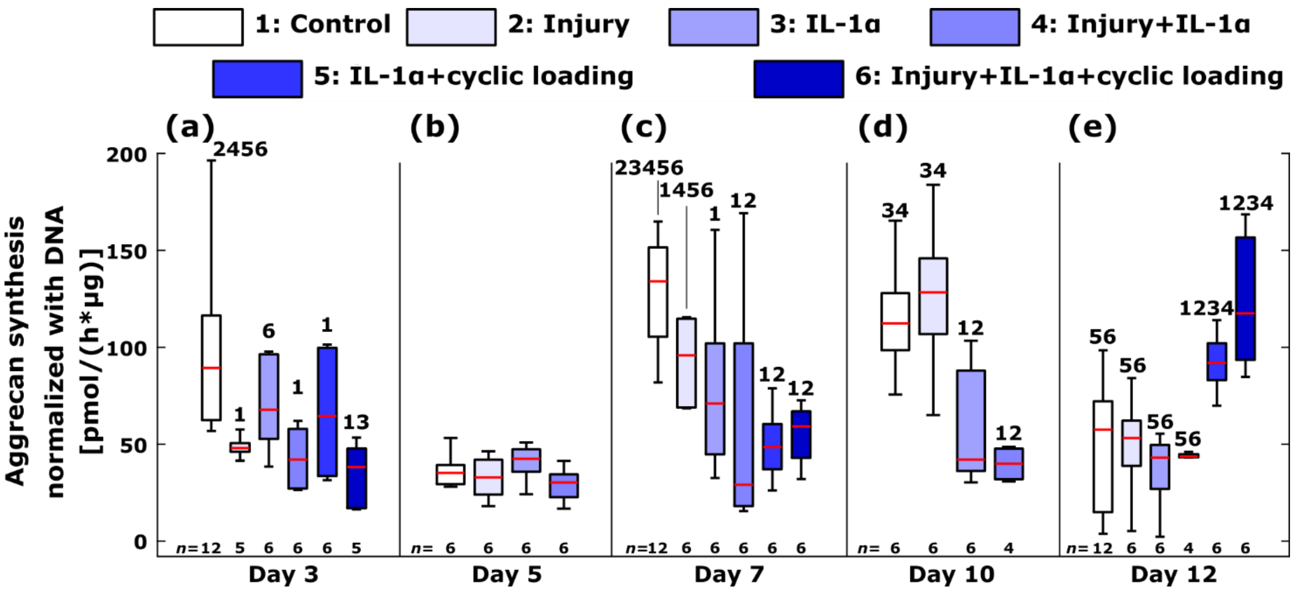


**Fig. S10. Aggrecan biosynthesis ($^{35}$S incorporation) normalized with the amount of DNA left in the chondrocytes between different treatment groups at different time points.** Aggrecan biosynthesis rate dropped fast after introduction of mechanical injury or pro-inflammatory cytokines. However, cyclic loading started to show its beneficial effects on aggrecan biosynthesis rate within 12 days from the initiation of the treatment. The amount of plugs *n* designated for each group is shown on the bottom of each subpanel. The numbers above boxplots show statistically significant difference in aggrecan biosynthesis rate compared to the indicated group(s) ($p < 0.05$; linear mixed effects model).

**Table S13. Aggrecan biosynthesis rate.** The results are shown as the estimated marginal mean values (pmol/h/μg) with standard error from the linear mixed effect models. *N* is the number of animals and *n* the total amount of plugs combined from those animals. The total amount of animals was 6. No cyclically loaded groups were terminated on days 5 and 10.

| | Day | | | | |
|---|---|---|---|---|---|
| **Group** | **3** | **5** | **7** | **10** | **12** |
| **CTRL** | **108.82<br>(11.41)<br>*N* = 4<br>*n* = 12** | **55.20<br>(14.62)<br>*N* = 2<br>*n* = 6** | **125.14<br>(11.25)<br>*N* = 4<br>*n* = 12** | **97.06<br>(32.06)<br>*N* = 2<br>*n* = 6** | **38.76<br>(11.40)<br>*N* = 4<br>*n* = 12** |
| **INJ** | **64.04<br>(15.38)<br>*N* = 2<br>*n* = 5** | **51.14<br>(14.62)<br>*N* = 2<br>*n* = 6** | **88.01<br>(14.04)<br>*N* = 2<br>*n* = 6** | **108.10<br>(14.61)<br>*N* = 2<br>*n* = 6** | **31.20<br>(14.61)<br>*N* = 2<br>*n* = 6** |

| | | | | | |
|---|---|---|---|---|---|
| **IL** | 88.58<br>(14.62)<br>*N* = 2<br>*n* = 6 | 59.01<br>(14.26)<br>*N* = 2<br>*n* = 6 | 75.11<br>(14.04)<br>*N* = 2<br>*n* = 6 | 38.74<br>(14.61)<br>*N* = 2<br>*n* = 6 | 18.48<br>(14.61)<br>*N* = 2<br>*n* = 6 |
| **INJ-IL** | 61.37<br>(14.62)<br>*N* = 2<br>*n* = 6 | 47.74<br>(14.62)<br>*N* = 2<br>*n* = 6 | 55.27<br>(14.04)<br>*N* = 2<br>*n* = 6 | 22.57<br>(16.64)<br>*N* = 2<br>*n* = 4 | 25.97<br>(16.56)<br>*N* = 2<br>*n* = 4 |
| **IL-CL** | 63.30<br>(14.57)<br>*N* = 2<br>*n* = 6 | N.A. | 47.40<br>(14.57)<br>*N* = 2<br>*n* = 6 | N.A. | 89.62<br>(14.57)<br>*N* = 2<br>*n* = 6 |
| **INJ-IL-CL** | 31.35<br>(15.40)<br>*N* = 2<br>*n* = 5 | N.A. | 52.96<br>(14.57)<br>*N* = 2<br>*n* = 6 | N.A. | 120.50<br>(14.57)<br>*N* = 2<br>*n* = 6 |

**Table S14. Descriptive statistics of the differences in estimated marginal means of aggrecan biosynthesis between different time points within groups.** The aggrecan biosynthesis rates are shown in the Table S13. The comparisons are shown between days 3, 5, 7, 10, and 12 with 95% confidence intervals and the associated *p*-value. The *p*-values smaller than 0.05 and 0.01 are shown with light yellow and light red, respectively. The difference is shown as the mean difference in the aggrecan biosynthesis (pmol/h/µg) in the specified group in row minus the group in column. No cyclically loaded groups were terminated on days 5 and 10.

| | **CTRL Day** | | | | **INJ Day** | | | |
|---|---|---|---|---|---|---|---|---|
| **Day** | **3** | **5** | **7** | **10** | **3** | **5** | **7** | **10** |
| **5** | -53.62<br>(-81.96; -25.28)<br>$p < .001$ | | | | -12.90<br>(-45.45; 19.65)<br>$p = .435$ | | | |
| **7** | 16.32<br>(-6.23; 38.88)<br>$p = .155$ | 69.94<br>(40.28; 99.60)<br>$p < .001$ | | | 23.96<br>(-10.05; 57.98)<br>$p = .166$ | 36.86<br>(4.08; 69.65)<br>$p = .028$ | | |
| **10** | -11.75<br>(-43.09; 19.58)<br>$p = .459$ | 41.86<br>(5.61; 78.12)<br>$p = .024$ | -28.08<br>(-57.77; 1.62)<br>$p = .064$ | | 44.06<br>(6.65; 81.47)<br>$p = .021$ | 56.96<br>(20.70; 93.21)<br>$p = .002$ | 20.10<br>(-12.74; 52.93)<br>$p = .228$ | |
| **12** | -70.06<br>(-93.89; -46.22)<br>$p < .001$ | -16.44<br>(-47.76; 14.88)<br>$p = .301$ | -86.38<br>(-108.95; -63.81)<br>$p < .001$ | -58.30<br>(-86.65; -29.96)<br>$p < .001$ | -32.84<br>(-70.25; 4.57)<br>$p = .085$ | -19.94<br>(-56.20; 16.31)<br>$p = .279$ | -56.80<br>(-89.64; -23.97)<br>$p = .001$ | -76.90<br>(-107.88; -45.92)<br>$p < .001$ |
| | **IL Day** | | | | **INJ-IL Day** | | | |
| **Day** | **3** | **5** | **7** | **10** | **3** | **5** | **7** | **10** |
| **5** | -29.58<br>(-60.56; 1.41)<br>$p = .061$ | | | | -13.63<br>(-44.61; 17.35)<br>$p = .386$ | | | |
| **7** | -13.47<br>(-46.26; 19.31)<br>$p = .418$ | 16.10<br>(-16.69; 48.89)<br>$p = .333$ | | | -6.11<br>(-38.89; 26.68)<br>$p = .713$ | 7.53<br>(-25.26; 40.31)<br>$p = .651$ | | |
| **10** | -49.84<br>(-86.10; -13.59)<br>$p = .007$ | -20.27<br>(-56.52; 15.97)<br>$p = .271$ | -36.37<br>(-69.21; -3.53)<br>$p = .030$ | | -38.80<br>(-78.15; 0.55)<br>$p = .053$ | -25.17<br>(-64.52; 14.18)<br>$p = .208$ | -32.70<br>(-68.69; 3.30)<br>$p = .075$ | |
| **12** | -70.10<br>(-106.36; -33.85)<br>$p < .001$ | -40.53<br>(-76.78; -4.27)<br>$p = .029$ | -56.63<br>(-89.47; -23.79)<br>$p = .001$ | -20.26<br>(-51.24; 10.72)<br>$p = .198$ | -35.40<br>(-74.47; 3.67)<br>$p = .075$ | -21.77<br>(-60.84; 17.30)<br>$p = .272$ | -29.30<br>(-65.14; 6.55)<br>$p = .108$ | 3.40<br>(-34.72; 41.53)<br>$p = .860$ |
| | **IL-CL Day** | | | | **INJ-IL-CL Day** | | | |
| **Day** | **3** | **5** | **7** | **10** | **3** | **5** | **7** | **10** |

| | | | | | | | | |
|---|---|---|---|---|---|---|---|---|
| **5** | N.A. | | | | N.A. | | | |
| **7** | -15.90 (-46.88; 15.08) $p = .312$ | N.A. | | | 21.62 (-10.93; 54.16) $p = .191$ | N.A. | | |
| **10** | N.A. | N.A. | N.A. | | N.A. | N.A. | N.A. | |
| **12** | 26.32 (-4.66; 57.30) $p = .095$ | N.A. | 42.22 (11.24; 73.20) $p = .008$ | N.A. | 89.16 (56.61; 121.70) $p < .001$ | N.A. | 67.54 (36.56; 98.52) $p < .001$ | N.A. |

**Table S15. Descriptive statistics of the differences in estimated marginal means of aggrecan biosynthesis between different groups at several time points.** The aggrecan biosynthesis rates are shown in the Table S13. The comparisons are shown between different groups on days 3, 5, 7, 10, and 12 with 95% confidence intervals and the associated *p*-value. The *p*-values smaller than 0.05 and 0.01 are shown with light yellow and light red, respectively. The difference is shown as the mean difference in the aggrecan biosynthesis (pmol/h/µg) in the specified group in row minus the group in column. No cyclically loaded groups were terminated on days 5 and 10.

| | Day 3 Group | | | | | Day 5 Group | | | | |
|---|---|---|---|---|---|---|---|---|---|---|
| **Group** | **CTRL** | **INJ** | **IL** | **INJ-IL** | **IL-CL** | **CTRL** | **INJ** | **IL** | **INJ-IL** | **IL-CL** |
| **INJ** | -44.77 (-74.74; -14.81) $p = .004$ | | | | | -4.06 (-35.04; 26.93) $p = .796$ | | | | |
| **IL** | -20.23 (-48.57; 8.11) $p = .160$ | 24.54 (-8.01; 57.09) $p = .138$ | | | | 3.81 (-27.17; 34.79) $p = .808$ | 7.86 (-23.12; 38.85) $p = .616$ | | | |
| **INJ-IL** | -47.44 (-75.78; -19.10) $p = .001$ | -2.67 (-35.22; 29.88) $p = .871$ | -27.21 (-58.19; 3.77) $p = .085$ | | | -7.46 (-38.44; 23.53) $p = .635$ | -3.40 (-34.38; 27.58) $p = .828$ | -11.27 (-42.25; 19.72) $p = .473$ | | |
| **IL-CL** | -45.52 (-73.86; -17.18) $p = .002$ | -0.75 (-37.97; 36.48) $p = .968$ | -25.29 (-61.29; 10.72) $p = .167$ | 1.92 (-34.08; 37.93) $p = .916$ | | N.A. | N.A. | N.A. | N.A. | |
| **INJ-IL-CL** | -77.47 (-107.52; -47.42) $p < .001$ | -32.70 (-71.20; 5.81) $p = .095$ | -57.24 (-94.60; -19.87) $p = .003$ | -30.03 (-67.39; 7.34) $p = .114$ | -31.95 (-64.50; 0.60) $p = .054$ | N.A. | N.A. | N.A. | N.A. | N.A. |

| | Day 7 Group | | | | | Day 10 Group | | | | |
|---|---|---|---|---|---|---|---|---|---|---|
| **Group** | **CTRL** | **INJ** | **IL** | **INJ-IL** | **IL-CL** | **CTRL** | **INJ** | **IL** | **INJ-IL** | **IL-CL** |
| **INJ** | -37.13 (-65.15; -9.12) $p = .010$ | | | | | 11.04 (-19.94; 42.02) $p = .482$ | | | | |
| **IL** | -50.03 (-78.05; -22.01) $p = .001$ | -12.90 (-43.88; 18.08) $p = .412$ | | | | -58.32 (-89.30; -27.34) $p < .001$ | -69.36 (-100.34; -38.38) $p < .001$ | | | |
| **INJ-IL** | -69.87 (-97.89; -41.85) $p < .001$ | -32.74 (-63.72; -1.76) $p = .039$ | -19.84 (-50.82; 11.14) $p = .207$ | | | -74.49 (-109.43; -39.55) $p < .001$ | -85.53 (-120.47; -50.59) $p < .001$ | -16.17 (-51.11; 18.78) $p = .362$ | | |
| **IL-CL** | -77.74 (-105.76; -49.72) $p < .001$ | -40.61 (-75.56; -5.66) $p = .023$ | -27.71 (-62.66; 7.24) $p = .119$ | -7.87 (-42.82; 27.08) $p = .657$ | | N.A. | N.A. | N.A. | N.A. | |

| | | | | | | | | | | |
|---|---|---|---|---|---|---|---|---|---|---|
| **INJ-IL-CL** | **-72.18 (-100.19; -44.16) *p* < .001** | **-35.05 (-69.99; -0.10) *p* = .049** | **-22.15 (-57.10; 12.80) *p* = .212** | **-2.31 (-37.26; 32.64) *p* = .896** | **5.56 (-25.42; 36.54) *p* = .723** | **N.A.** | **N.A.** | **N.A.** | **N.A.** | **N.A.** |

| | **Day 12** | | | | |
|---|---|---|---|---|---|
| **Group** | **CTRL** | **INJ** | **IL** | **INJ-IL** | **IL-CL** |
| **INJ** | **-7.56 (-35.90; 20.79) *p* = .599** | | | | |
| **IL** | **-20.28 (-48.63; 8.07) *p* = .159** | **-12.72 (-43.70; 18.26) *p* = .418** | | | |
| **INJ-IL** | **-12.79 (-45.16; 19.59) *p* = .436** | **-5.23 (-40.16; 29.70) *p* = .768** | **7.49 (-27.44; 42.43) *p* = .672** | | |
| **IL-CL** | **50.86 (22.51; 79.20) *p* = .001** | **58.41 (22.40; 94.43) *p* = .002** | **71.14 (35.12; 107.15) *p* < .001** | **63.64 (24.59; 102.70) *p* = .002** | |
| **INJ-IL-CL** | **81.74 (53.40; 110.09) *p* < .001** | **89.30 (53.29; 125.31) *p* < .001** | **102.02 (66.01; 138.04) *p* < .001** | **94.53 (55.48; 133.58) *p* < .001** | **30.89 (-0.10; 61.87) *p* = .051** |
